# Multiple Quality Control Checkpoints Safeguard Small Nuclear RNA Biogenesis and Prevent Assembly of Aberrant Spliceosomes

**DOI:** 10.1101/2025.11.21.689835

**Authors:** Tiantai Ma, Claire Huntington, Zhuoyi Song, Rea M. Lardelli, Eric L. Van Nostrand, Jens Lykke-Andersen

## Abstract

Defective small nuclear (sn)RNAs are produced from hundreds of human snRNA pseudogenes and mutant snRNA genes associated with human developmental disorders. Machineries that prevent defective snRNAs from disrupting pre-mRNA splicing remain poorly defined. Here, we identify multiple checkpoints in snRNA biogenesis monitored by quality control machineries that subject defective snRNAs to degradation and prevent their assembly into spliceosomes. We show that variant U1 snRNAs produced from human pseudogenes, some at rates approaching the canonical snRNAs, are impaired in 3’ cleavage and targeted for degradation by the NEXT-exosome while failures in subsequent protein assembly steps promote NEXT-exosome- or Terminal Uridylyl Transferase 4/7-mediated degradation. These pathways also repress mutant snRNAs associated with human developmental disorders. Impeding snRNA quality control causes formation of aberrant spliceosomes and altered pre-mRNA splicing. These findings define checkpoints in snRNA biogenesis that safeguard pre-mRNA splicing and represent potential therapeutic targets for human disorders associated with snRNA mutations.

## Introduction

The integrity of the pre-mRNA splicing machinery is essential for eukaryotic gene expression and aberrant splicing is associated with scores of human disorders^1,2^. Mutations in small nuclear (sn)RNAs, the central components of the spliceosome responsible for pre-mRNA splicing, have been associated with human developmental disorders including neurodevelopmental diseases^3–10^. Yet, hundreds of snRNA pseudogenes exist in the human genome with the potential to produce snRNA variants containing nucleotide substitutions in residues critical for pre-mRNA splicing^11–16^. The capacity of these snRNA variants to impact pre-mRNA splicing and the machineries that may exist to prevent their adverse effects remain poorly defined.

U1 snRNA variants in particular appear to bew highly transcribed and capable of impacting pre-mRNA splicing^14–16^. Previous studies presented evidence for association of RNA polymerase II (Pol II) with U1 snRNA pseudogenes at levels that rival those of canonical U1 snRNA genes^16^. Moreover, U1 snRNA variants have been observed to alter spliceosome 5’ splice site specificity when exogenously expressed in cells^15^. Yet, U1 snRNA variants accumulate at levels at least three orders of magnitude below canonical snRNAs^14^ and those U1 snRNA variants whose stability have been tested were observed to be rapidly degraded^14,17^. Thus, quality control machineries must exist that detect and degrade transcribed snRNA variants.

Defective snRNAs can also be produced from canonical snRNA genes through mutation. Recent studies have identified mutations in canonical U4-2, U2-2P and U5 snRNA genes as major factors in neurodevelopmental disorders (NDD)^3,5,8^. Moreover, mutations in minor class U4atac snRNA have been linked to developmental disorders^4,6,9,18^ and mutations in minor class U12 snRNA has been shown to cause early-onset cerebellar ataxia and CDAGS syndrome^10,19^. Additionally, U1 and U2 snRNAs have been reported to be mutated in multiple cancers^20,21^. Whether these disease-associated mutant snRNAs are targeted by quality control pathways remains unknown.

The biogenesis of Pol II-transcribed snRNAs, which include all spliceosomal snRNAs except U6 and U6atac, is a multi-step process that involves both nuclear and cytoplasmic events^17,22^. Nascently transcribed snRNAs undergo co-transcriptional 7-methyl guanosine (m7G) capping followed by 3’ end cleavage by the Integrator complex^23,24^, which leaves a short snRNA 3’ end tail. The newly transcribed snRNAs are exported to the cytoplasm by export factor PHAX via interaction with the nuclear Cap-Binding Complex (CBC) bound to the m7G cap^25^. After entering the cytoplasm, snRNAs assemble with the Sm complex and are then subjected to further cap methylation to form the Tri-Methyl Guanosine (TMG) cap^26–30^. Sm complex-assembled, TMG capped snRNAs are subsequently re-imported into the nucleus^31,32^ and subjected to nucleotide modifications by sno/scaRNAs^33^. During these nuclear and cytoplasmic biogenesis steps, snRNAs also assemble with snRNA-specific proteins, which in the case of U1 snRNA includes U1A, U1C, and U1-70K^33–35^. One of the last steps in snRNA biogenesis involves trimming of the snRNA 3’ end tail by exonuclease TOE1^22^. Thus, nascent snRNAs for the majority of their biogenesis exist with 3’ tails that may expose them to 3’ end degradation machineries.

Here, we identify multiple checkpoints in snRNA biogenesis that are monitored by quality control pathways, which subject defective snRNA variants produced from pseudogenes and snRNA mutants associated with human disorders for degradation from the 3’ end. These checkpoints monitor accurate 3’ end cleavage and snRNA protein complex assembly and subject defective snRNAs to degradation by the NEXT-exosome complex or Terminal Uridylyl Transferases (TUTs) 4/7. Inactivation of the NEXT-exosome complex causes stabilization of U1 snRNA variants, assembly of U1 snRNA variants with spliceosomes, and alterations in pre-mRNA splicing. Thus, multiple snRNA biogenesis checkpoints ensure proper spliceosome assembly by preventing the accumulation of defective snRNAs and may contribute to developmental disorders by repressing the expression of disease-associated mutant snRNAs.

## Results

### Highly transcribed human snRNA variants are degraded during early biogenesis

The human genome harbors snRNA pseudogenes across all chromosomes (Figure 1A). To facilitate genome-wide analyses of the fates of snRNA variants produced from pseudogenes, we created a high-throughput sequencing alignment pipeline designed to minimize faulty mapping of highly abundant reads arising from canonical snRNA genes to snRNA pseudogenes (Figure 1B). Leveraging this pipeline, we first asked which snRNA pseudogenes are actively transcribed. Analysis of a Pol II ChIP-seq dataset from human Hela cells^36^ showed evidence - consistent with previous studies^16^ - of high levels of Pol II association with a subset of snRNA pseudogenes, the majority of which were U1 snRNA pseudogenes (Figure 1C). Furthermore, analysis of ENCODE ChIP-seq datasets for components of the snRNA gene-specific SNAPc transcription complex in human HepG2 cells similarly showed high association with a subset of snRNA pseudogenes (Supplementary Figures S1A-C)^37,38^. To ask whether Pol II-associated snRNA pseudogenes could be observed to be transcribed, and if the resulting snRNA variants accumulate to the steady state, we next applied our pipelines to a HeLa cell nascent RNA-seq dataset^39^. Indeed, a large number of snRNA variants can be observed in the nascent RNA population but are seen at much reduced levels in the steady-state population (Figure 1D and Supplementary Table S2). This suggests that a subset of snRNA variants are highly transcribed but are rapidly degraded, consistent with previous observations for individually tested variants^14,17^.

**Figure 1:**
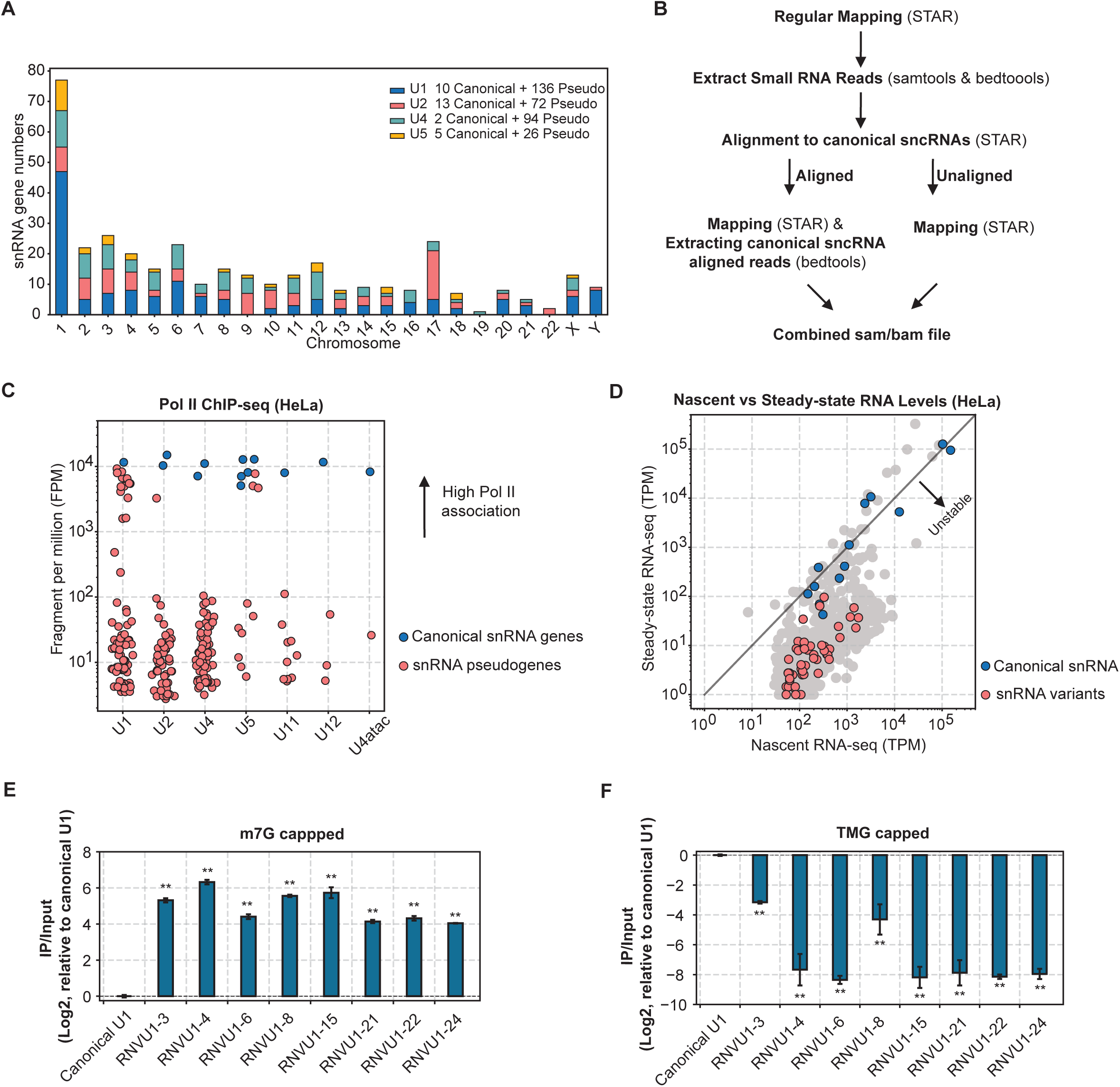
A large number of human snRNA variants are transcribed but degraded during early biogenesis. (A) Major class Pol II snRNA gene distribution across human chromosomes. The number of snRNA genes (GENCODE V33 annotation) is plotted for each chromosome. (B) Diagram of the 3-pass STAR alignment pipeline to detect snRNA variant reads from high-throughput sequencing datasets.(C) Scatter plot of Pol II ChIP-seq reads, represented as fragments per million (FPM), for individual Pol II snRNA canonical genes (blue) and pseudogenes (red). Dataset from^36^. See Supplementary Table S2 for canonical versus pseudogene definitions. (D) Scatter plot of HeLa cell nascent versus steady-state sncRNA levels by RNA seq, measured as transcripts per million reads (TPM). Canonical snRNAs are labeled in blue, snRNA variants in red, and other small non-coding RNAs in grey. Dataset from^39^. (E) Relative levels of m7G-capped U1 snRNA variants compared to canonical U1 snRNA, measured by m7G-cap IP followed by RT-qPCR and normalized to ratios in input RNA samples. P-values were calculated by Student’s two-tailed t-test comparing variant IP/input values with those for canonical U1 snRNA (**: P<0.01). (F) Same as in (E) but monitoring TMG cap-IP.

To test whether snRNA variants undergo degradation during biogenesis, prior to their full maturation, we took advantage of the fact that the nascent snRNA m7G cap undergoes further methylation in late biogenesis to form a TMG cap^22^. Immunoprecipitation followed by RT-qPCR assays revealed that U1 snRNA variants are highly enriched for m7G-capped and de-enriched for TMG-capped species relative to canonical U1 snRNA (Figure 1E and 1F). Thus, snRNA variants are degraded during biogenesis prior to maturation.

### SnRNA variants are inefficiently 3’ end cleaved

An important step in early snRNA processing is co-transcriptional 3’ end cleavage by the Integrator complex, which is thought directed by a conserved 3’ box sequence occurring immediately downstream of canonical snRNA genes (Figure 2A)^24,40–42^. All U1 snRNA variants indicated by Pol-II ChIP as highly transcribed deviate from the consensus by one or more nucleotides in the 3’ box sequences (Figure 2B). Canonical snRNAs that are produced with 3’ end extensions due to failure in 3’ end cleavage have been observed to undergo degradation^43^. We therefore wondered whether U1 snRNA variants are unstable due to inefficient Integrator-mediated 3’ cleavage. Indeed, metagene analyses of Pol II ChIP-seq datasets from human HeLa cells^36^ revealed increased association of Pol II with downstream regions of U1 snRNA pseudogenes as compared to canonical U1 snRNA genes (Figure 2C). This pattern could be observed also for individual U1 snRNA pseudogenes (Figure 2D). A similar analysis of a HeLa cell ChIP-seq dataset for INTS11^36^, the endonuclease component of the Integrator complex, similarly shows increased association with U1 snRNA pseudogenes into the downstream regions as compared to the canonical genes (Figures 2E and 2F), suggesting delayed recruitment or release of INTS11 during transcription. Furthermore, analyzing an INTS11 depletion RNA-seq dataset from HeLa cells^36^ revealed a significant increase in the accumulation of canonical snRNAs with 3’ end extensions upon INTS11 depletion, whereas snRNA variants were mostly unaffected (Figure 2G), consistent with the idea that snRNA variants are poorly 3’ end cleaved by Integrator. To test whether U1 snRNA variants are indeed produced with 3’ end extensions, we analyzed chromatin-associated and nuclear RNA-seq datasets from HeLa cells^44^. We found that an increased fraction of U1 snRNA variants harbor 3’ end extensions in comparison to canonical U1 snRNA as observed in metagene analyses (Figure 2H) as well as in analyses of individual snRNA variants (Figure 2I). Taken together, our observations show that U1 snRNA pseudogenes are defective in 3’ end cleavage and produce snRNA variants with extended 3’ ends.

**Figure 2:**
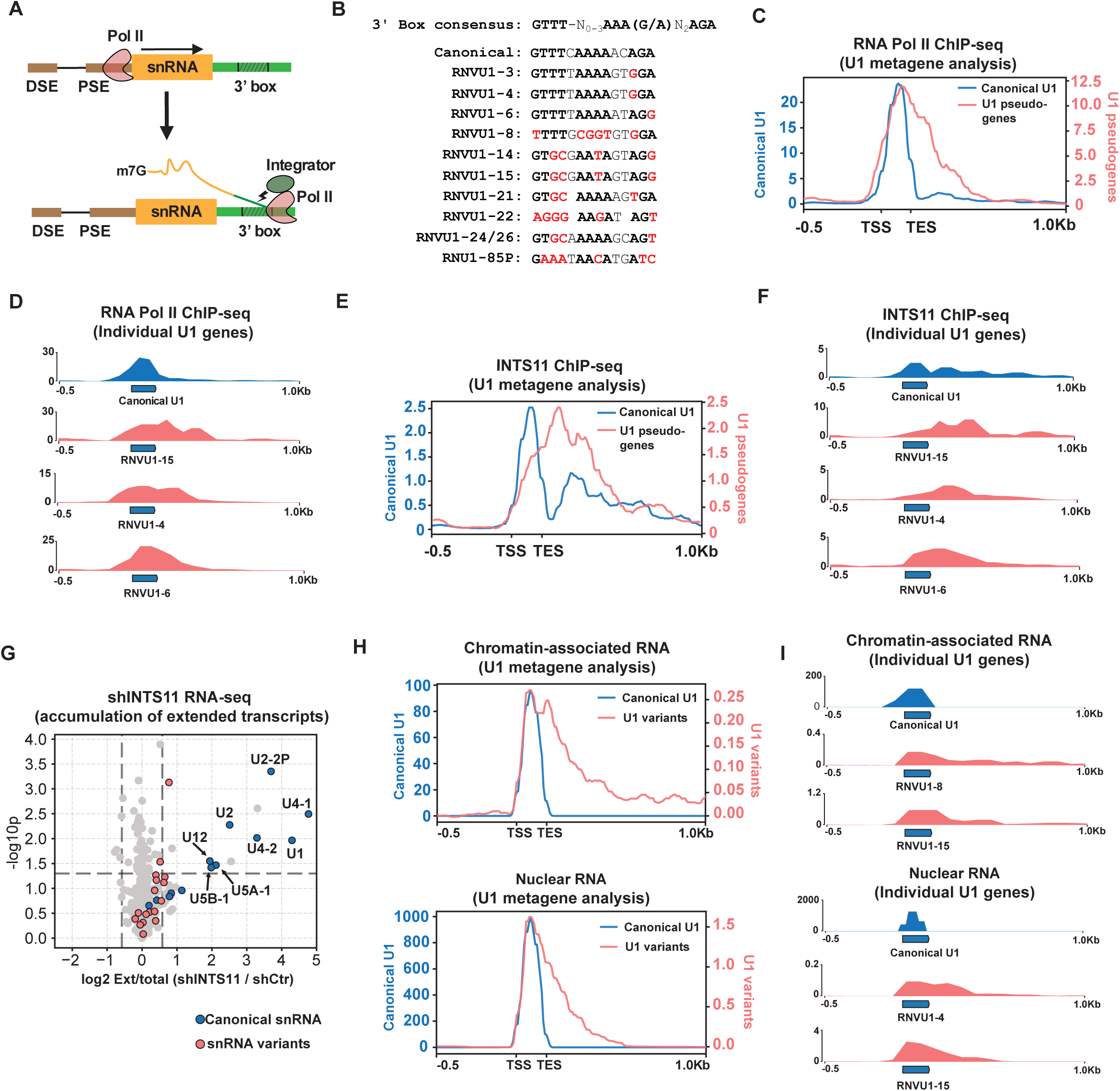
snRNA variants are inefficiently 3’ end cleaved. (A) Schematic of snRNA transcription by RNA Pol II. snRNA gene elements, DSE, PSE, and 3’ box, are shown at their relative snRNA gene position. Integrator cleavage is shown as a lighting symbol. (B) Alignment of 3’ box sequences of canonical U1 snRNA genes and U1 snRNA pseudogenes. Conserved 3’ box nucleotides are shown in bold black. Nucleotides in snRNA pseudogenes that do not match consensus are shown in bold red. (C) Metagene plot showing average distribution of HeLa cell Pol II ChIP-seq reads for canonical U1 genes and all transcribed U1 pseudogenes. Dataset from^36^. Values on the left represent average BPM (Bins per million mapped reads) per canonical U1 gene and on the right represent average U1 pseudogene signal. Signals from 0.5Kb upstream of the transcription start site (TSS) to 1Kb downstream of the transcription end site (TES) are plotted. (D) Individual signal tracks for average canonical U1 genes and select U1 pseudogenes from the Pol II ChIP-seq dataset in (C). (E) Metagene plot as in (B) but plotting INTS11 ChIP-seq signals from^36^ (F) Signal track as in (D) monitoring INTS11 ChIP-seq. (G) Volcano plot showing accumulation of 3’ end extended sncRNAs in INTS11 depletion (shINTS11) relative to control (shCtr) conditions, measured as the log2 fold ratios in shINTS11 condition over control conditions of reads 1 to 500 nucleotides downstream of annotated 3’ ends over the sum of gene body and downstream TPMs. P-values represent three individual biological repeats calculated by Student’s two-tailed t-test. Canonical snRNAs are labeled in blue, snRNA variants in red, and other sncRNAs in grey. (H) Metagene plots for chromatin-associated (upper panel) and nuclear (lower panel) RNA read distributions as described in (C). RNA-seq datasets are from^44^. (I) Individual track plots for chromatin-associated and nuclear RNA reads for select U1 snRNA genes as described in (D).

### The NEXT-exosome complex degrades 3’ end extended snRNA variants

The NEXT-exosome complex is known to degrade canonical snRNAs with 3’ end extensions^43^ (Figure 3A). To test whether snRNA variants are subjected to degradation by the NEXT-exosome, we first applied our snRNA variant analysis pipeline to an RNA-seq dataset from HeLa cells in which the MTR4 RNA helicase component of the NEXT complex was depleted^45,46^. We found that all U1 snRNA variants observed in the dataset are significantly upregulated upon MTR4 depletion (Figure 3B). Analysis of read coverage for individual transcripts demonstrated that for canonical U1 snRNAs the vast majority of reads correspond to the snRNA body, but 3’ end extended molecules can be observed to accumulate upon MTR4 depletion. By contrast, U1 snRNA variants show similar levels of upregulation of both the gene body and 3’ end extensions (Figure 3C) consistent with poor Integrator cleavage and MTR4-dependent degradation of the 3’ end extended molecules.

**Figure 3:**
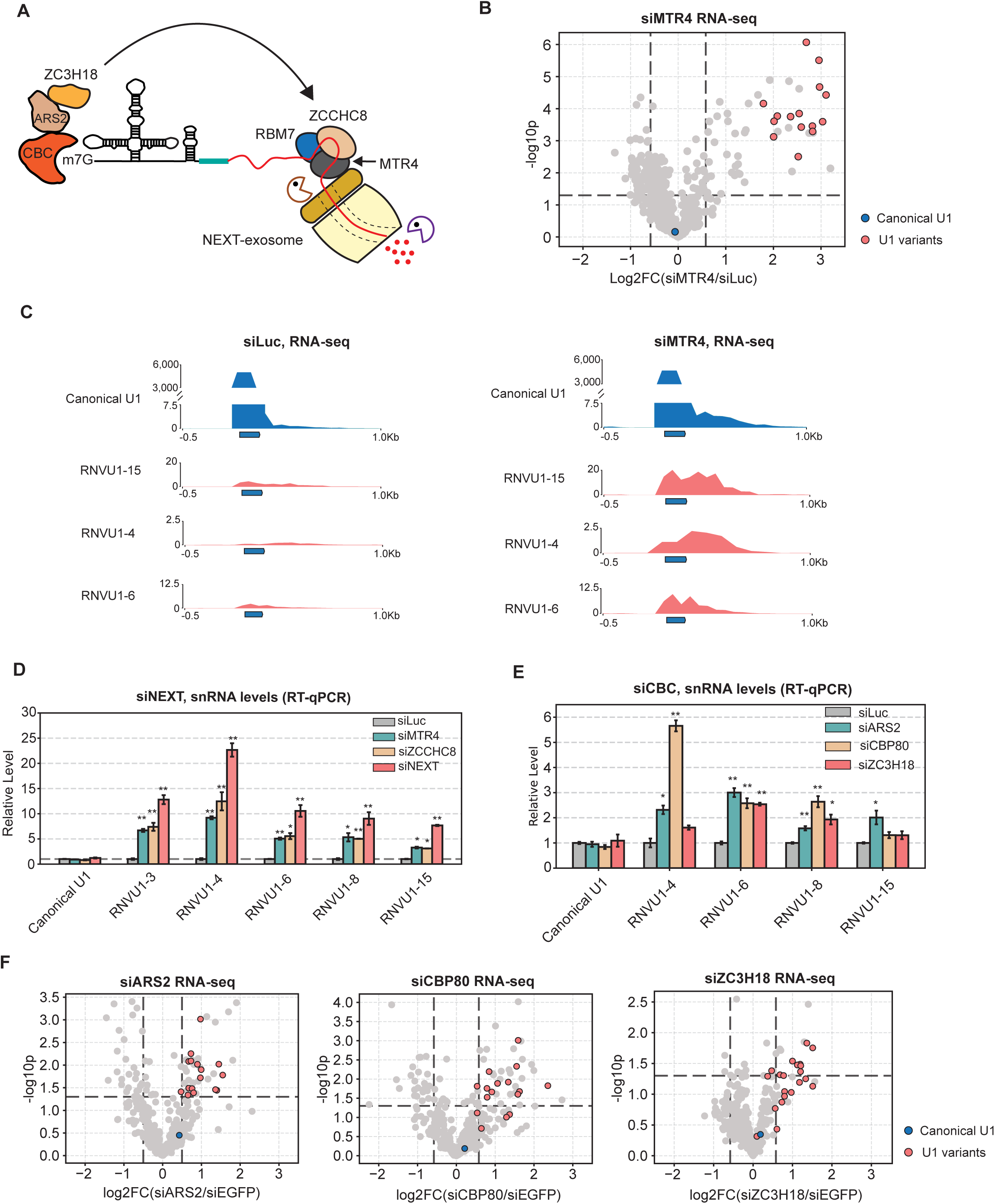
The NEXT-exosome complex degrades 3’ end extended snRNA variants. (A) Schematic of the NEXT-exosome complex degrading 3’ extended snRNAs stimulated by the CBC. (B) Volcano plot showing the log2 fold ratios in sncRNA levels (RNA-seq; TPM) in HeLa cells depleted for MTR4 (siMTR4) over control (siLuc). P-values were calculated from three individual biological repeats by Student’s two-tailed t-test. Canonical snRNAs are labeled in blue, snRNA variants in red, and other small non-coding RNAs in grey. Dataset is from^45,46^. (C) RNA seq signal tracks for canonical U1 and select U1 variants in control (siLuc, left) and MTR4-depleted (siMTR4, right) conditions from the dataset in (B). Signals from 0.5Kb upstream of the transcription start site (TSS) to 1Kb downstream of the transcription end site (TES) are plotted. (D) Relative levels of canonical U1 and U1 variant snRNAs upon depletion of NEXT-exosome components (siNEXT represents co-depletion of MTR4 and ZCCHC8) quantified by RT-qPCR and normalized to siLuc control conditions. P-values were calculated by two-tailed Student’s t-tests (*: p<0.05, **: p<0.01). (E) Relative levels of canonical U1 and U1 variant snRNAs as in (D), but in CBP80, ARS2 or ZC3H18 depletion conditions. (F) Volcano plots as in (B) monitoring changes in sncRNA levels upon ARS2 depletion (left), CBP80 depletion (middle), and ZC3H18 depletion (right) relative to control depletion conditions. Datasets are from^52^.

MTR4 is the central helicase component of multiple exosome adapter complexes, including the NEXT, TRAMP and PAXT complexes^47^. To test which of these complexes is responsible for the observed U1 snRNA variant degradation we used RT-qPCR assays to monitor the impact of depletion of components specific to each of the MTR4 complexes on canonical U1 snRNA and U1 snRNA variant levels (Supplementary Figures S2A and S2B). We found that all tested U1 snRNA variants are significantly upregulated upon depletion of NEXT complex subunit ZCCHC8 (Figure 3D), whereas depletion of ZCCHC7 and ZFC3H1 components of TRAMP and PAXT complexes, respectively, showed only minor or no impact on accumulation of snRNA variants (Supplementary Figures S2C and S2D).

The NEXT-exosome complex is known to associate with the nuclear Cap Binding Complex (CBC) via ARS2 and ZC3H18^48–51^ (Figure 3A). To test whether this association is important for snRNA variant degradation, we tested the impact of knocking down ARS2, CBP80, and ZC3H18 on U1 snRNA variant accumulation. All tested U1 snRNA variants saw increased accumulation upon depletion of one or more of these components, although some variants were more sensitive than others (Figure 3E). Furthermore, analyses of published RNA seq datasets^52^ using our snRNA variant pipeline revealed accumulation of snRNA variants upon depletion of each of these factors (Figure 3F), consistent with the idea that NEXT-exosome mediated degradation of snRNA variants is functionally linked to the CBC (Figure 3A). By contrast, depletion of the CBC-binding snRNA export factor PHAX or of the mRNA-specific CBC-binding protein NCBP3 did not significantly impact the accumulation of snRNA variants (Supplementary Figures S3E and S3F). Collectively, our analyses show that snRNA variants are defective in 3’ end cleavage and targeted for degradation by the NEXT-exosome complex, stimulated by the CBC.

### Defective U1-70K binding triggers U1 snRNA degradation by the NEXT-exosome independent of a 3’ end cleavage defect

We have previously observed U1 snRNA variants with 3’ tails similar in length to Integrator-cleaved canonical snRNAs^17,22^, suggesting that a subset of snRNA variant transcription events undergo normal 3’ end cleavage. To monitor the fate of variant snRNAs with normal 3’ end cleavage, we tested the impact of replacing the native 50 bp downstream sequence of an exogenously expressed barcoded RNVU1-15 variant^22^ with the 50 bp downstream sequence of the canonical U1 snRNA RNU1-2 gene, including the 3’ box sequence thought to direct 3’ end cleavage (Figure 4A). Using targeted RNA sequencing, we found that this modification, as expected, resulted in a significant decrease in RNVU1-15 3’ extended transcripts (Figure 4B) and a significant increase in the overall accumulation of RNVU1-15 RNA (Figure 4C). However, this 3’ end-modified RNVU1-15 variant continued to accumulate at lower levels than a co-expressed canonical U1 snRNA (Figure 4C) and remained sensitive to NEXT complex depletion (Figure 4D), suggesting that there may be features of the RNVU1-15 variant other than 3’ end cleavage efficiency that trigger instability.

**Figure 4:**
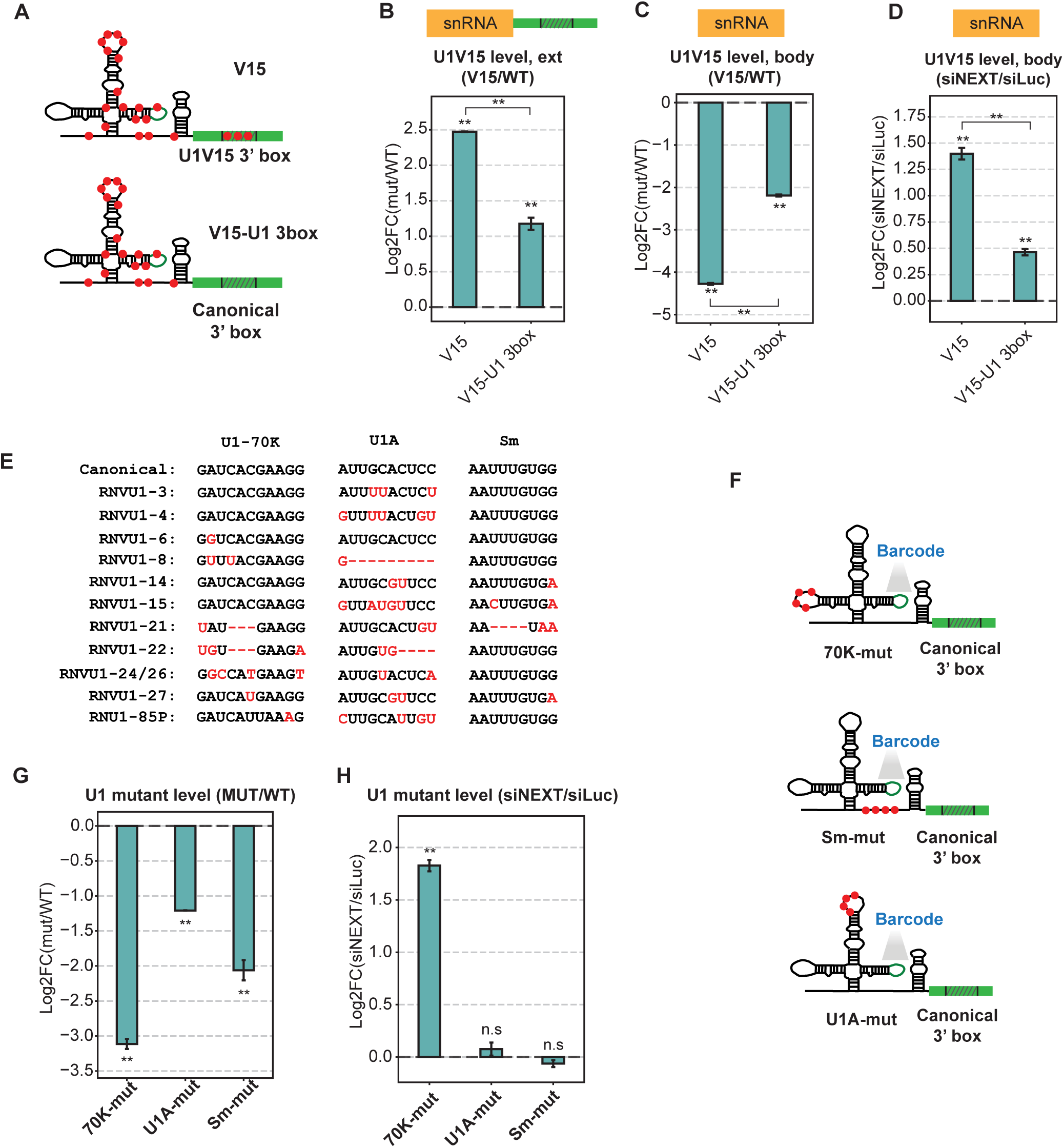
U1-70K binding site mutation triggers U1 snRNA degradation by the NEXT-exosome independent of a 3’ end cleavage defect. (A) Schematics of V15 (upper) and V15-U1 3box (lower) RNAs. Variant nucleotides are shown as red dots. (B) Log2 fold change in accumulation of 3’ extended V15 RNAs relative to co-expressed canonical U1 (WT), quantified by targeted RNA-seq. P-values comparing level of individual extended V15 RNAs with extended WT are labeled above each bar and the p-value for comparison of the two V15 RNAs is labeled between the bars; Student’s two-tailed t-test. (**: p<0.01). (C) Log2 fold difference in total accumulation of V15 RNAs (mature plus extended) relative to co-expressed canonical U1 (WT), quantified by targeted RNA-seq. P-values are calculated as in (B). (D) Log2 fold change in accumulation of V15 RNAs in NEXT depletion (siNEXT) over control (siLuc) conditions, normalized to co-expressed canonical U1 (WT), quantified by targeted RNA-seq. P-values are calculated as in (B). (E) Sequence alignment of binding sites for U1-70K, U1A, and the Sm complex in canonical U1 and highly expressed U1 variant snRNAs. Variant nucleotides are shown in red. (F) Schematics of 70K-mut, U1A-mut, and Sm-mut U1 snRNAs with mutations indicated as red dots and a barcode sequence in stem loop 3 indicated as a green line. (G) Log2 fold difference in accumulation of U1 mutants relative to co-expressed U1 WT, quantified by targeted RNA-seq. P-values were calculated as in (B). (H) Log2 fold change in U1 mutant snRNA accumulation in NEXT depletion (siNEXT) over control (siLuc) conditions quantified by targeted RNA-seq and normalized to co-expressed U1 WT. P-values were calculated as in (B) (n.s.: p>0.5).

U1 snRNA variants also harbor nucleotide variations in protein binding sites (Figure 4E). To test whether protein assembly impacts U1 snRNA accumulation, we generated barcoded canonical U1 snRNA reporters with mutations previously established to abolish interaction with specific U1 snRNP proteins^28,53,54^ (Figure 4F). Using targeted sequencing, we found that all tested protein assembly-deficient U1 snRNA mutants accumulate at significantly lower levels than a co-expressed unmutated U1 snRNA (Figure 4G). This reduction in U1 snRNA accumulation is not caused by impaired 3’ end cleavage because all mutant constructs contained canonical U1 snRNA downstream sequences and produced similar levels of 3’ end extended transcripts as the co-expressed unmutated U1 snRNA (Supplementary Figure S3). Depletion of the NEXT complex led to significant upregulation of the U1-70K binding-deficient mutant U1 snRNA, but had no effect on the other U1 snRNA mutants (Figure 4H). Thus, the NEXT-exosome complex targets not only U1 snRNAs that fail to undergo correct 3’ cleavage but also a U1 snRNA that fails to assemble with U1-70K.

### U1A and Sm complex binding site mutations trigger TUT4/7-mediated U1 snRNA degradation

Previous studies have shown that Sm-mutant U1 snRNAs are unstable^55,56^ and accumulate post-transcriptional oligouridine tails added by cytoplasmic terminal-uridylyl transferases (TUT) 4 and 7^22,56,57^. We therefore next tested whether TUT4/7 depletion impacts the accumulation of U1 snRNA protein-binding mutants (Supplementary Figures S4A and S4B). Indeed, mutant U1 snRNAs defective in Sm assembly and, to a less extent, U1A assembly are significantly upregulated upon TUT4/7 depletion, whereas the U1-70K assembly mutant saw no impact (Figure 5A). Similarly, U1 snRNA engineered to contain RNVU1-15-specific variations in Sm or U1A binding sites saw increased accumulation upon TUT4/7 depletion but not upon NEXT complex depletion (Supplementary Figures S4C-E). Using RT-qPCR assays to test the accumulation of select endogenous snRNA variants, RNVU1-15 and two U2 snRNA variants, RNVU2-17P and RNVU2-63P, are significantly upregulated following TUT4/7 depletion (Figure 5B). These upregulated snRNA variants all harbor variations in the Sm binding site (Figure 5C).

**Figure 5:**
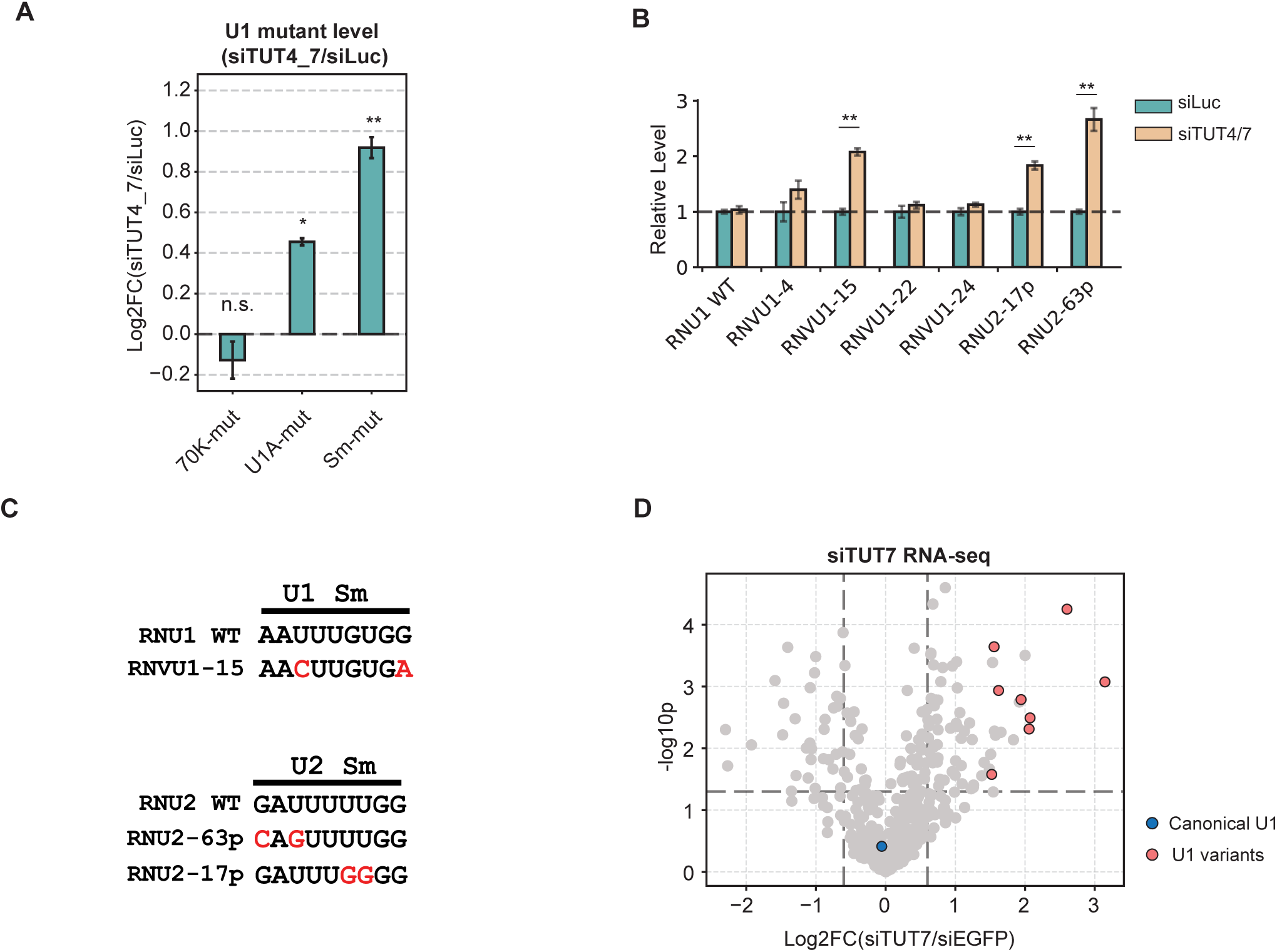
U1A and Sm complex binding site mutations trigger TUT4/7-mediated U1 snRNA degradation. (A) Log2 fold change in accumulation of U1 snRNA mutants in TUT4/7-depleted (siTUT4_7) over control (siLuc) conditions, quantified by targeted RNA-seq. P-values are calculated through two-tailed Student’s t-test (n.s.: p>0.05, *:p<0.05, **: p<0.01). (B) Relative accumulation of canonical U1 and snRNA variants under control (siLuc) or TUT4/7-depleted (siTUT4/7) conditions quantified by RT-qPCR. P-values are calculated by two-tailed Student’s t-test. (C) Sequence alignment of Sm binding sites of snRNA variants upregulated under siTUT4/7 conditions. Nucleotide variations are shown in red. (D) Volcano plot showing the log2 fold change in sncRNA accumulation in TUT7-depleted (siTUT7) over control (siEGFP) conditions monitored by RNA-seq. P-values were calculated from three individual biological repeats by Student’s two-tailed t-test. Canonical snRNAs are labeled in blue, snRNA variants in red, and other small non-coding RNAs in grey. Dataset is from^52^.

We further analyzed TUT7 and TUT4 depletion RNA-seq datasets from a human PA-1 ovarian cancer cell line^58^. While RNVU1-15 was the only U1 snRNA variant significantly upregulated upon TUT4 depletion (Supplementary Figure S4F), all detected U1 snRNA variants saw significant upregulation upon depletion of TUT7 (Figure 5D). This is consistent with the fact that nearly all U1 snRNA variants harbor variations in U1A and/or Sm binding sites (Figure 4D) and suggests that there may be some differences in which snRNA variants are targeted by TUT4 versus TUT7. Thus, defective U1 snRNAs are monitored by both NEXT-exosome and TUT4/7 degradation machineries, which recognize distinct snRNA features.

### SnRNA variants assemble with spliceosomes upon NEXT complex depletion

Since the identified quality control checkpoints eliminate snRNA variants during snRNA biogenesis, we wondered whether they are critical for preventing snRNA variants from assembling into the spliceosome. To address this question, we used enhanced (e)CLIP^59,60^ to monitor the association of snRNA variants with spliceosome components SNRNP200 (BRR2) and PRPF8 (PRP8) in the absence or presence of NEXT complex depletion (Figure 6A and Supplementary Figures S5A-B). These specific spliceosome factors were selected because their antibodies have been previously verified for eCLIP^59,60^ and, as components of the U4/U6.U5 tri-snRNP^61^, they are known to associate with the U1 snRNP only after initiation of the pre-mRNA splicing process. Since neither PRPF8 nor SNRNP200 interact directly with U1 snRNA, we used formaldehyde-rather than UV-crosslinking to ensure capture of the entire spliceosome complex. This has been shown to be effective for enriching U1 snRNA in previous studies^17^, and we indeed observe enrichment of several snRNAs including U1, as well as of pre-mRNA introns over exons, by this method (Supplementary Figures S5C-E).

**Figure 6.**
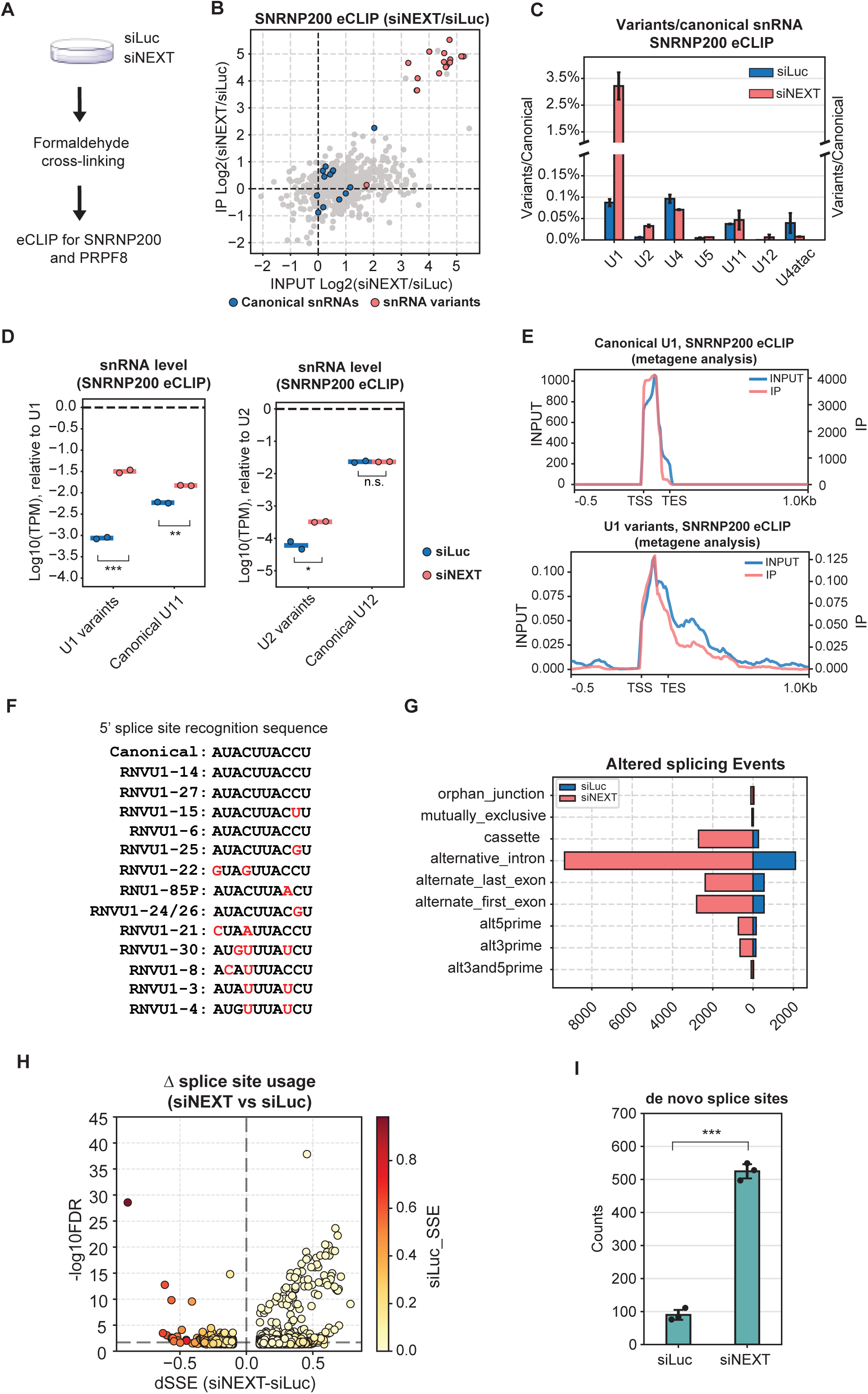
SnRNA variants assemble with spliceosomes upon NEXT-exosome depletion. (A) Schematic of eCLIP conditions. (B) Scatter plot of sncRNA enrichment in SNRNP200 eCLIP, showing ratios of reads (measured as TPM) in NEXT depletion (siNEXT) over control (siLuc) conditions in Input versus IP fractions. Canonical snRNAs are labeled in blue, U1 variants in red, and other sncRNAs in grey. (C) Bar plot showing ratios of aggregate snRNA variant levels relative to corresponding canonical snRNA levels in SNRNP200 eCLIP IP samples in siLuc (blue) and siNEXT (red) conditions. (D) U1 variant (in aggregate) and canonical U11 snRNA levels relative to canonical U1 snRNA levels (measured as TPM) in SNRNP200 eCLIP IP samples in siLuc (blue) and siNEXT (red) conditions. P-values are calculated through two-tailed Student’s t-test (n.s.: p>0.05, *: p<0.05 **: p<0.01, ***: p<0.001). (E) Metagene plots showing read distributions for canonical U1 snRNAs (upper panel) and U1 variants (lower panel) in SNRNP200 eCLIP input (blue) and IP (red) conditions. Signals from 0.5Kb upstream of the transcription start site (TSS) to 1Kb downstream of the transcription end site (TES) are plotted. (F) Sequence alignment of 5’ splice site recognition sequences in canonical U1 and U1 variants. (G) Altered splicing events in eCLIP input samples, quantified by MAJIQ^80^, in siLuc (blue) and siNEXT (red) conditions compared to human genome annotation. (H) Volcano plot showing differential splice site (dSSE) usage between siNEXT and siLuc conditions in eCLIP input samples quantified by SpliSER^62^. Dots represent individual splice sites colored according to fractional splice site usage in the siLuc condition (scale on the right). Only splice sites with dSSE values >0.05 or <-0.05 are shown. (I) Quantification of numbers of *de novo* splice sites observed in siLuc or siNEXT conditions in (H). Splice sites with -log10FDR >2 and SSE values in opposing conditions of less than 0.05 are considered as *de novo* splice sites. P-value was calculated by two-tailed Student’s t-test (***: p<0.001).

We found that for a majority of observed U1 snRNA variants, NEXT complex depletion resulted in a 10- to 50-fold increase in their association with SNRNP200 and PRPF8 as compared with control depletion conditions (Figures 6B and Supplementary Figure S5E). Indeed, U1 snRNA variants were, in aggregate, observed to increase from less than 0.1% of the total SNRNP200- and PRPF8-associated U1 snRNA population in control conditions to 1.5 to 3% (Figure 6C and Supplementary Figures S5G-H) upon NEXT-complex depletion, reaching levels higher than that observed for U11 snRNA of the minor spliceosome (Figures 6D and Supplementary Figure S5I). These observations suggest that NEXT-exosome depletion allows the assembly of spliceosomes containing variant U1 snRNAs at levels that exceed that of the minor spliceosome.

The level of U2 snRNA variants associated with SNRNP200 and PRPF8 also increased upon NEXT depletion (Figures 6C-D and Supplementary Figures S5H-I), mostly due to one U2 variant, RNU2-63P (Supplementary Figures S5G). By contrast, the fraction of variants of other major and minor class snRNAs remained low and unchanged upon NEXT complex depletion, consistent with low variant expression among other snRNA classes (Figures 1C-D). Evaluating snRNA read coverage in the eCLIP assays suggested that spliceosome assembly is selective towards variant U1 snRNAs that have undergone some amount of 3’ end trimming but also accommodates some 3’ end extended molecules (Figures 6E and Supplementary Figure S5J).

The formation of U1 snRNA variant spliceosomes upon NEXT complex depletion raised the possibility of impacts on pre-mRNA splicing. Indeed, a subset of the most abundant snRNA variants contain nucleotide variations in 5’ splice site recognition sequences (Figure 6F) and previous studies have demonstrated that exogenous expression of U1 snRNA variants impacts 5’ splice site usage^15^. Analyzing reads from our eCLIP input RNA dataset, we observed a greater number of altered splicing events under NEXT depletion conditions compared to control conditions (Figure 6G). To more specifically monitor the impact on the usage of individual splice sites, we utilized the SpliSER pipeline^62^. This revealed hundreds of splice sites that saw significantly increased usage upon NEXT complex depletion, far outweighing in number the splice sites that moved in the opposite direction, and the vast majority of which were *de novo* sites, rarely observed in control conditions (Figures 6H-I). Examination of representative genes seeing accumulation of mRNAs with *de novo* splice sites upon NEXT complex depletion confirms accumulation of alternative mRNA isoforms (Supplementary Figure S6A). While mRNA stabilization rather than altered pre-mRNA splicing may underlie the accumulation of a subset of mRNAs with *de novo* splicing events, a majority were associated with genes that did not see increased transcript levels upon NEXT complex depletion (Supplementary Figures S6B-D). Thus, consistent with previous observations of altered pre-mRNA splicing upon exogenous expression of U1 snRNA variants^15^, this suggests direct alterations in splice site usage as a consequence of aberrant spliceosome assembly. Taken together, these observations suggest that the NEXT-exosome prevents the assembly of U1 snRNA variants into spliceosomes and alterations in pre-mRNA splicing.

### SnRNA quality control pathways target mutant snRNAs associated with human developmental disorders

Mutations in canonical snRNA genes have been associated with developmental disorders, including neurodevelopmental disease (NDD)^3,5,8^. We considered whether disease-associated mutant snRNAs, similarly to snRNA variants, may be targeted by quality control pathways. Previous studies have observed degradation by the nuclear exosome of a mutant U12 snRNA associated with Early Onset Cerebellar Ataxia^7,10^ and reduced accumulation of a subset of mutant U4atac snRNAs associated with developmental disorders^4^. We used exogenous expression assays to test the impact on snRNA accumulation of NDD-associated 64insU, A65G and 77insU mutations in U4-2 snRNA as well as of G48A, G51A, G55A, and U120G mutations in U4atac snRNA associated with human developmental disorders^4,63,64^ (Figure 7A). When transiently expressed in HeLa cells, each of these mutant snRNAs accumulated at significantly lower levels than their co-expressed unmutated snRNA controls (Figure 7B), although the impact of the 77insU mutation on U4-2 snRNA accumulation was minor. The LWS-associated U120G U4atac mutant, which was the most significantly downregulated snRNA mutant tested, could be observed to accumulate with an unprocessed 3’ end extension and a post-transcriptional oligo-U tail (Supplementary Figure S7A-C).

**Figure 7.**
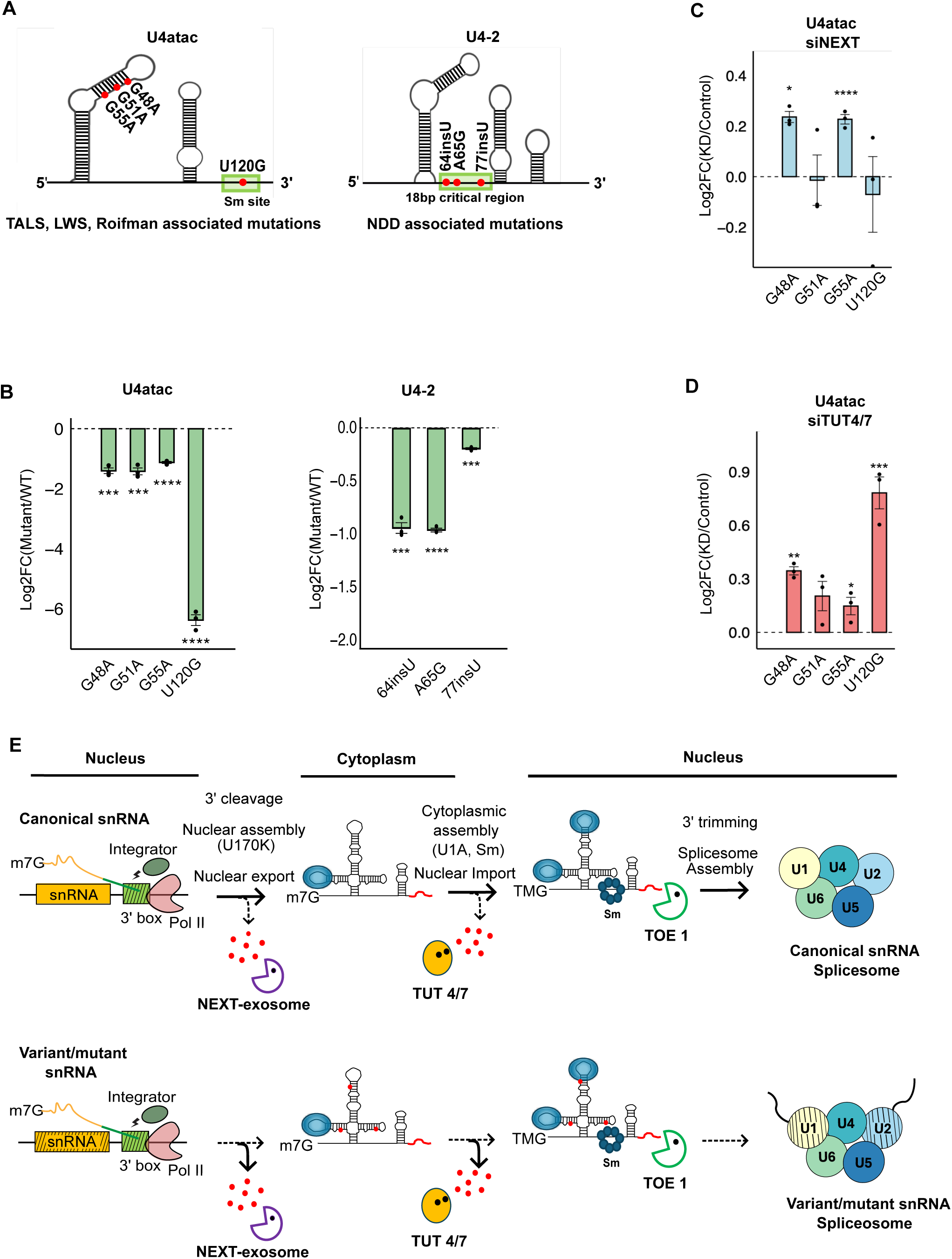
SnRNA quality control pathways target mutant snRNAs associated with human developmental disorders. (A) Schematic of U4atac and U4-2 snRNAs highlighting tested disease-associated mutations. (B) Accumulation of U4atac and U4-2 mutant snRNAs relative to co-expressed WT snRNAs, quantified by targeted RNA-seq. P-values were calculated using two-tailed Student’s t-test and labeled beneath bars (*: p<0.1, **: p<0.05, ***: p<0.01, ****: p<0.001). (C) Accumulation of U4atac mutant snRNAs in siNEXT over siLuc control conditions, quantified by targeted RNA-seq and normalized to co-expressed U4atac WT snRNA. (D) Accumulation of U4atac mutant snRNAs in siTUT4/7 over siLuc control conditions, quantified as in (C). (E) Model showing checkpoints in snRNA biogenesis monitored by quality control identified in this study and the assembly of snRNA variants/mutants into variant snRNA spliceosomes in the absence of those pathways.

To determine if the quality control pathways that target U1 snRNA variants also contribute to the observed downregulation of disease-associated mutant snRNAs, we tested the impact of depletion of the NEXT complex and TUT4/7 on the accumulation of U4atac mutant snRNAs. Depletion of the NEXT complex led to upregulation of the RFMN-associated U4atac G48A mutant and the MOPD1-associated U4atac G55A mutant, whereas the levels of U4atac G51A and U120G mutants remained unchanged (Figure 7C). Upon depletion of TUT4/7, all tested U4atac mutants accumulated at increased levels with three of four reaching statistical significance. Collectively, these observations demonstrate the targeting of disease-associated mutant snRNAs by the same quality control pathways that target snRNA variants, with individual mutants exhibiting different sensitivities to different quality control factors.

## Discussion

The human genome harbors hundreds of snRNA pseudogenes a subset of which are transcribed at rates approaching those of canonical snRNA genes (Figure 1), but the pathways that ensure that defective snRNA variants produced from pseudogenes do not interfere with normal pre-mRNA splicing and the impact of such pathways on mutant snRNAs associated with human developmental disorders have remained poorly defined. In this study, we identify multiple quality control checkpoints that monitor proper snRNA biogenesis and repress the production of defective variant snRNAs and mutant snRNAs associated with human diseases. One of these checkpoints monitors co-transcriptional 3’ end cleavage by the Integrator complex, which we demonstrate is defective for all tested U1 snRNA variants (Figure 2), and subjects those that are aberrantly cleaved to degradation by the NEXT-exosome complex stimulated by the m7G cap-associated CBC complex (Figure 3). Additional checkpoints monitor snRNA assembly with proteins and, in the case of U1 snRNAs, target those defective in assembly with U1-70K for degradation by the NEXT-exosome complex (Figure 4) and those with U1A and Sm complex assembly defects for TUT4/7-mediated degradation (Figure 5). These snRNA biogenesis checkpoints serve to prevent the formation of aberrant spliceosomes as evidenced by the assembly of U1 and U2 snRNA variants with spliceosomes and *de novo* splice site usage upon depletion of the NEXT-exosome (Figure 6). The snRNA quality control checkpoints also target mutant snRNAs associated with human developmental disorders as evidenced by the negative impact of disease-associated mutations on U4-2 and U4atac snRNA accumulation and the targeting of U4atac snRNA mutants by the NEXT-exosome complex and/or TUT4/7 (Figure 7). These findings uncover quality control checkpoints that monitor snRNA biogenesis and prevent the formation of aberrant spliceosomes containing defective snRNA variants and mutant snRNAs associated with human disorders.

Defects in U1 snRNA 3’ end cleavage and U1-70K assembly both trigger degradation by the NEXT-exosome complex, but how does the NEXT-exosome monitor each of these processes? The observations that U1 snRNA variants are stabilized upon depletion of the CBC subunit CBP80 and of factors that link the CBC to the NEXT-exosome complex demonstrate that U1 snRNA variant degradation is stimulated by the m7G cap. Previous reports have presented evidence for a kinetic competition between the snRNA nuclear export factor PHAX and the NEXT-exosome-associated protein Z3CH18 for binding to the CBC^52,65,66^. Moreover, the nuclear export of m7G-capped small RNAs by PHAX has been reported to be less efficient the longer the RNA^67^. Taken together, these findings suggest that NEXT-exosome mediated degradation of snRNA variants occurs in competition with nuclear export, such that 3’ end extended snRNA variants experience delayed nuclear export, affording the time for NEXT-exosome-mediated degradation via kinetic competition with PHAX for CBC binding (Figure 7E). This raises the possibility that U1-70K assembly is monitored by a similar competition between nuclear export and degradation by the exosome. Indeed, U1-70K assembly is known to promote U1 snRNA Sm complex assembly^35^ and 3’ end trimming by TOE1^22^, consistent with a step critical for early biogenesis.

Other U1 snRNA biogenesis checkpoints monitor U1A and Sm complex assembly and depend on TUT4/7-mediated degradation. The known localization of TUT4/7^57^ and of Sm complex assembly^68^ in the cytoplasm suggests that this is a cytoplasmic snRNA biogenesis checkpoint. Consistent with this idea, previous studies have implicated cytoplasmic factors, 3’ exonuclease DIS3L2 and the Lsm1-7-decapping complex, in degrading U1 snRNAs containing mutations in the Sm binding site^55,56^. The stage in biogenesis at which U1A assembles with U1 snRNA is less well understood. Our observations suggest that U1A assembly occurs either prior to nuclear export or upon localization in the cytoplasm, either of which is compatible with a cytoplasmic quality control step. Sm complex assembly is known to promote TMG capping and subsequent nuclear import of snRNAs, and thus, in analogy to the competition between nuclear export and NEXT-exosome degradation, snRNA nuclear import may act in kinetic competition with TUT4/7-mediated snRNA degradation in the cytoplasm (Figure 7E).

Our observation that U4-2 and U4atac snRNA mutations associated with developmental disorders cause snRNA downregulation and that tested mutant U4atac snRNAs are targeted by the same degradation machineries that target U1 snRNA variants raise the possibility that snRNA degradation contributes to developmental disorders associated with snRNA mutations. Similarly, mutant U12 snRNA associated with Early Onset Cerebellar Ataxia^7^ was observed targeted by the nuclear exosome^10^. In addition to the U4-2 snRNA mutations tested here, mutations in U2-2P and U5 snRNAs have also been implicated in NDD^3,5,8^. Recent studies have presented evidence that U4-2 and U5 snRNA mutations lead to NDD at least in part due to specific splicing defects^3^. An important question for future study is to what degree downregulation via quality control checkpoints identified here contribute to the dysfunction of these disease-associated mutant snRNAs and whether snRNA stabilization could be a productive therapeutic avenue.

Depletion of the NEXT-exosome complex led to a dramatic increase in the association of U1 snRNA variants with spliceosomal factors SNRP200 and PRPF8 at levels above U11 snRNA of the minor spliceosome. Moreover, consistent with a subset of the most highly expressed U1 snRNA variants possessing nucleotide variation in 5’ splice site recognition sequences and with tested U1 snRNA variants having been observed to alter 5’ splice site usage when exogenously expressed^15^, we observed *de novo* usage of hundreds of splice sites correlating with U1 snRNA variant spliceosome accumulation upon NEXT-exosome depletion. This suggests that the quality control checkpoints identified here play critical roles in safeguarding the pre-mRNA splicing process by preventing the formation of variant spliceosomes that would otherwise accumulate at levels beyond the minor spliceosome. These findings also raise the possibility that snRNA variants may become activated in certain tissues or conditions to generate specialized spliceosomes, which, in analogy with the minor spliceosome, may drive specialized splicing programs.

Our findings identify the snRNA 3’ end as a hub for quality control checkpoints that monitor snRNA biogenesis. We have previously observed that snRNAs undergo a final 3’ end trimming step by the snRNA processing enzyme TOE1 during late stages of biogenesis, after Sm complex assembly and TMG capping^22^ (Figure 7E). The maintenance of an snRNA 3’ tail until the final stages of biogenesis likely allows for the multiple 3’ end-mediated quality control checkpoints identified here, highlighting the importance of competing maturation- and decay-promoting exonucleases in the quality control of small non-coding RNA biogenesis. Our findings highlight what may well be an important general principle in RNA biogenesis whereby individual processing steps are monitored by quality control checkpoints to ensure that only fully functional RNPs are allowed to accumulate in the cell, thereby preventing adverse effects of canonical RNAs that experience failures during processing and of the thousands of small non-coding RNA pseudogenes that exist in the human genome (Figure 7E).

## Acknowledgements

We thank Amy Pasquinelli, Cody Ocheltree, Megan Dowdle, and Kalodiah Toma for useful discussions of the manuscript. This work was supported by NIH grant R35GM118069 to J.L-A and R35HG011909 to E.L.V.N. R.M.L. was the recipient of a National Research Service Award Postdoctoral Fellowship (NIH F32 GM106706) and was a San Diego IRACDA Fellow (NIH K12 GM06852). We acknowledge the UCSD Stem Cell Genomics and Microscopy Core for use of their Illumina Miseq instrument. This publication includes data generated at the UC San Diego IGM Genomics Center utilizing an Illumina NovaSeq X Plus that was purchased with funding from a National Institutes of Health SIG grant (#S10 OD026929).

## Author contributions

Conceptualization: T.M., R.M.L., J.L-A.; Data curation: T.M., C.H., Z.S., R.M.L.; Formal analysis: T.M., C.H., Z.S.; Investigation: T.M., C.H., Z.S.; Software: T.M., C.H., Z.S.; Supervision: R.M.L., E.L.V.N., J.L-A.; Visualization: T.M., C.H.; Writing – original draft: T.M, J.L-A.; Writing – review & editing: all authors.

## STAR METHODS

### Key resources table

| REAGENT or RESOURCE | SOURCE | IDENTIFIER |
| --- | --- | --- |
| Antibodies |  |  |
| Anti-7-methylguanosine (m7G)-Cap mAb | MBL | RN016M |
| Anti-2,2,7-trimethylguanosine (m3G/TMG) mAb | MBL | RN019M |
| Mouse IgG2b Isotype Control | ThermoFisher | Cat #02-6300 |
| Rabbit anti-PRPF8 Antibody | Bethyl Lab | Cat #A303-921A |
| Rabbit anti-SNRNP200 Antibody | Bethyl Lab | Cat #A303-453A |
| Mouse anti-ZCCHC8 antibody | Abcam | Cat #ab68739 |
| Rabbit anti-MTR4 antibody | Abcam | Cat #ab70551 |
| Rabbit anti-ZCCHC6 (TUT7) | Sigma | Cat #HPA020615 |
| Rabbit polyclonal anti-β-actin | Cell Signaling | Cat #4967S |
| Donkey anti-Rabbit IgG (H+L) Cross-Adsorbed Secondary Antibody, HRP | ThermoFisher | Cat #31485 |
| Goat anti-Rabbit IgG (H+L) Secondary Antibody, HRP | ThermoFisher | Cat #31430 |
| Bacterial and virus strains |  |  |
| DH5-alpha competent *E.coli* strain | N/A | N/A |
| Chemicals, peptides, and recombinant proteins |  |  |
| siLentFect™ Lipid Reagent for RNAi | Bio-Rad | Cat #1703362 |
| Lipofectamine™ 2000 Transfection Reagent | ThermoFisher | Cat #11668027 |
| Turbo DNase | ThermoFisher | Cat #AM2239 |
| Proteinase K | NEB | Cat #M0681L |
| SuperScript™ III Reverse Transcriptase | ThermoFisher | Cat #18080044 |
| T4 Polynucleotide Kinase | NEB | Cat #P8107S |
| FastAP Alkaline Phosphatase | ThermoFisher | Cat #EF0651 |
| T4 RNA Ligase 1 | NEB | Cat #M0204L |
| T4 RNA Ligase 1 (ssRNA Ligase), High Concentration | NEB | Cat #M0437M |
| RNaseOUT™ Recombinant Ribonuclease Inhibitor | ThermoFisher | Cat #10777019 |
| SuperScript III reverse transcriptase | Invitrogen | Cat #18080044 |
| Protease Inhibitor Cocktail | ThermoFisher | Cat #A32953 |
| ExoSAP-IT PCR product cleanup reagent | Applied Biosystems | Cat #78201.1.ML |
| Critical commercial assays |  |  |
| RNA Clean & Concentrator-5 | Zymo | Cat #R1016 |
| PureLink™ HiPure Plasmid Maxiprep Kit | ThermoFisher | Cat #K210007 |
| SYBR™ Green Universal Master Mix | Applied Biosystems | Cat #4309155 |
| NEBuilder HiFi DNA Assembly cloning kit | NEB | Cat #E5520S |
| Gibson Assembly® Master Mix | NEB | Cat #E2611S |
| Deposited data |  |  |
| eCLIP | This study | GSE310123 |
| 3' end sequencing | This study | GSE310122 |
| Experimental models: Cell lines |  |  |
| HeLa S3 | N/A | N/A |
| Oligonucleotides |  |  |
| qRT-PCR oligonucleotides | See Table S1 | N/A |
| siRNA oligonucleotides | See Table S1 | N/A |
| Recombinant DNA |  |  |
| pU1-Barcoded U1WT | Ma et al.<sup>22</sup> | N/A |
| pU1-70K-mut | Ma et al.<sup>22</sup> | N/A |
| pU1-U1A-mut | Ma et al.<sup>22</sup> | N/A |
| pU1-Sm-mut | Ma et al.<sup>22</sup> | N/A |
| pU1-v15 | Ma et al.<sup>22</sup> | N/A |
| pU1-v15-U1A | Ma et al.<sup>22</sup> | N/A |
| pU1-v15-Sm | Ma et al.<sup>22</sup> | N/A |
| pU1-v15-U1 3 box | This study | N/A |
| pU4atac-U4atac-WT | This study | N/A |
| pU4atac-U4atac-G48A | This study | N/A |
| pU4atac-U4atac-G51A | This study | N/A |
| pU4atac-U4atac-G55A | This study | N/A |
| pU4atac-U4atac-U120G | This study | N/A |
| pU4-U4-2-WT | This study | N/A |
| pU4-U4-2-64insU | This study | N/A |
| pU4-U4-2-A65G | This study | N/A |
| pU4-U4-2-77insU | This study | N/A |
| Software and algorithms |  |  |
| Python (3.10.13/2.7.18) | N/A | N/A |
| STAR (v2.7.11b) | Github | https://anaconda.org/bioconda/star |
| Cutadapt (v5.1) | Conda | https://github.com/marcelm/cutadapt/ |
| UMI-tools (v1.1.6) | Github | https://github.com/CGATOxford/UMI-tools |
| deeptools (v3.5.6) | Github | https://github.com/deeptools/deepTools |
| MAJIQ (v2.5) | N/A | https://majiq.biociphers.org/ |
| SpliSER | Github | https://github.com/CraigIDent/SpliSER |
| CLIPper | Github | https://github.com/YeoLab/clipper |
| matplotlib (v3.8.3) | Github | https://github.com/matplotlib/matplotlib |
| ggplot2 (v3.3.5) | N/A | https://ggplot2.tidyverse.org/ |
| samtools (v1.22.1) | Conda | https://anaconda.org/bioconda/samtools |
| bedtools (v2.31.1) | Conda | https://anaconda.org/bioconda/bedtools |
| Tailer | Github | https://github.com/TimNicholsonShaw/tailer |
| Other |  |  |
| VAHTS DNA Clean Beads | Vazyme | Cat #N411-02 |
| Dynabeads™ Protein G | ThermoFisher | Cat #10003D |
| TRIzol™ Reagent | ThermoFisher | Cat #15596018 |

### Method Details

#### Plasmids

U1 snRNA expression plasmids pU1-barcoded U1WT, pU1-70K-mut, pU1-U1A-mut, pU1-Sm-mut, pU1-v15, pU1-v15-U1A, and pU1-v15-Sm were described previously^22^. Barcoded RNVU1-15 expression plasmid with a canonical U1 snRNA 3’ box (pU1-v15-U1 3 box) was generated by producing a PCR product of the barcoded RNVU1-15 sequence from pU1-v15 using V15_ligation_F and V15_overhang_R primers (Supplementary Table S1). Barcoded canonical U1 expression plasmid pU1-barcoded U1WT^22^ containing canonical RNU1-2 DSE, PSE, and 3’ box sequences was linearized by Polymerase Chain Reaction (PCR) using U1WT_bb_F and Flank_ligation_R primers (Supplementary Table S1) and ligated with the barcoded RNVU1-15 PCR product using Gibson Assembly (New England Biolabs) per manufacturer’s recommendation. U4-2 and U4atac barcoded expression vectors pU4-U4-2-WT and pU4atac-U4atac-WT were assembled using HiFi DNA assembly (New England Biolabs). The expression plasmid backbone was amplified from the pU1-barcoded U1WT^22^ plasmid as described above and gBlocks (IDT) containing human U4atac and U4-2 sequences with barcode mutations to allow distinction from the endogenous snRNAs (U4-2: C95G, G106C; U4atac: G94C, C102G) were assembled with the linearized plasmid; barcode mutations were designed to flip a stem-loop C-G base-pair in the human snRNAs to a G-C base-pair observed in other mammals. U4atac and U4-2 snRNA mutant expression plasmids (pU4atac-U4atac-G48A, pU4atac-U4atac-G51A, pU4atac-U4atac-G55A, pU4atac-U4atac-U120G, pU4-U4-2-64insU, pU4-U4-2-A65G, and pU4-U4-2-77insU) were generated in the same manner.

#### Cell culture, RNA interference, and plasmid transfections

All cells were maintained in Dulbecco’s Modified Eagle Medium (DMEM, Gibco) supplemented with 10% Fetal Bovine Serum (FBS, Gibco) and 1% penicillin/streptomycin (Gibco) at 37°C, 5% CO_2_. RNA interference was performed in HeLa cells with 20 nM small interfering (si)RNA targeting genes of interest (Supplementary Table S1), or luciferase as a control, using siLentFect (Bio-Rad) transfection reagent according to the manufacturer’s recommendations at 72 and 24 h before cell harvest. Plasmid transfections were performed using a total of 2 μg plasmid per 3.5-cm well plates using Lipofectamine 2000 (Life Technologies) transfection reagent according to the manufacturer’s recommendations at 48 h before harvest. U1 mutant plasmids were transfected in a ratio of 3:1 compared to U1WT plasmid. U4 and U4atac mutant plasmids were transfected in a ratio of 2:1 compared to U4 or U4atac WT plasmid, except for pU4atac-U4atac-U120G plasmid, which was transfected in a 4:1 ratio compared to U4atac WT plasmid.

#### Targeted RNA sequencing and analysis

RNA was isolated using TRIzol (Thermo Fisher) per the manufacturer’s recommendation. RNA adapters containing barcodes and 10- or 11-nt randomers (AG10/AG11; Supplementary Table S1) were ligated to the 3′ ends of 2.5 μg of extracted RNA in a 10-μl reaction containing 1 μl of 10xT4 RNA ligase buffer (500 mM Tris-HCl pH 7.5, 100 mM MgCl2, 10 mM DTT), 0.2 mg/mL bovine serum albumin (BSA), 2 μM AG10 or AG11, 1 mM ATP, 10 U of T4 RNA ligase (NEB), and 40 units of RNaseOUT (Thermo Fisher) for 16 h at 16°C. Ligated RNA was subsequently treated with DNase I (1 U/μL final concentration; Zymo research) in DNase buffer (10 mM Tris-HCl pH 7.5, 2.5 mM MgCl_2_, 0.5 mM CaCl_2_) at 25 °C for 15 min, and purified by RNA isolation and concentration using an RNA clean & concentrator-25 kit (Zymo Research). cDNA was generated using SuperScript III (SSRT III; Thermo Fisher) in a 20-μL reaction using 0.5 μM linker-specific primer AR17 (Supplementary Table S1). Targeted RNA 3′ end sequencing libraries were generated using gene-specific forward and reverse primers (Supplementary Table S1) and sequenced on an Illumina MiSeq platform as previously described^17,22^. FASTQ files were subjected to 3’ adaptor and PCR duplicate removal using custom python scripts. The reads were mapped to a custom reference using Tailer^69^ as previously described.

#### M7G/TMG cap immunoprecipitation

Total RNA from HeLa cells was isolated using TRIzol (Thermo Fisher) per the manufacturer’s recommendation. Genomic DNA was removed using a Turbo DNA-free kit (Thermo Fisher) per the manufacturer’s recommendation. 25 μl Dynabeads Protein G beads (Thermo Fisher) per sample were washed three times with NET-2 (10 mM Tris HCl pH 7.4, 150 mM NaCl, 0.1% Triton-X100), 1 x protease inhibitor cocktail (Thermo Fisher), and incubated with 5 μg of anti-m7G (MBL, RN016M), anti-TMG (MBL, RN019M), or mouse IgG (ThermoFisher, Cat #02-6300) antibody overnight at 4°C in 200 μl NET-2 with end-to-end rotation. 2 μg of the DNA-free RNA was added to 500 μl NET-2 and 1 μl RNaseOUT (40 U/µl; ThermoFisher) and incubated at 85°C for 5 mins and then on ice for 5 mins. After heating and cooling, an additional 1 μl RNaseOUT (40 U/µl) was added to the mix. 20 μl of antibody-liganded Protein G beads was added to each reaction followed by incubation at 4°C with end-to-end rotation for 2 hours. Beads were subsequently washed with 500 μl NET-2 eight times. After the final wash, 950 μl TRIzol was added to the beads. Immunoprecipitated RNA was subsequently isolated per the manufacturer’s recommendation.

#### Sequencing data analysis

FASTQ files were downloaded from NCBI Sequence Read Archive (SRA) or ENCODE Encyclopedia. Adaptors were trimmed using Cutadapt 5.1^70^. Adaptor-trimmed reads were mapped to the human genome (version hg38) using STAR 2.7.11b^71^ in a custom three-pass alignment pipeline described in Figure 1B. Briefly, sequencing reads were first aligned to a custom FASTA database of canonical small RNA genes, each including 50 base pair upstream and downstream sequences using (STAR --outFilterMultimapNma× 1000 --outFilterMultimapScoreRange 0 --outFilterMismatchNoverLmax 0.2 --outFilterMismatchNoverReadLmax 0.05 --clip5pNbases (if any) --clip3pNbases (if any) --alignIntronMin 9999999 -alignMatesGapMax 500 --alignEndsType EndToEnd --outReadsUnmapped Fastx). For ChIP-seq analysis, the first alignment step described above was skipped. Mapped (i.e. canonical sncRNA reads) and unmapped reads (i.e. reads that did not map to canonical sncRNAs) were each subsequently aligned to the full human genome using (STAR --outFilterMultimapNma× 1000 --outFilterMultimapScoreRange 0 --outFilterMismatchNoverLmax 0.025 --alignIntronMin 9999999 --alignMatesGapMax 2000 --clip5pNbases (if any) --clip3pNbases (if any) --alignEndsType EndToEnd) and, for the canonical sncRNA reads, those that did not map to canonical sncRNA genes were discarded using bedtools^72^. BAM/SAM files with aligned canonical and non-canonical small RNA reads were subsequently combined for downstream analysis. Mapped reads were counted using featureCounts^73^ or custom python scripts. Metagene plots were generated using deepTools^74^. Individual gene track was generated using Integrative Genomics Viewer (IGV) 2.17.4 or using pyGenomeTracks^75^.

#### eCLIP library preparation

eCLIP was adapted from the previously described protocol^76^, with modification of the crosslinking approach to stabilize indirect interactions within the spliceosome complex. Here, cells were crosslinked using 0.2% formaldehyde, followed by immunoprecipitation of spliceosomal complexes with antibodies against PRPF8 (#A303-921A, Bethyl Lab) or SNRNP200 (#A303-453A, Bethyl Lab). 2% of the lysate was saved as input and then denatured with 1X NuPage buffer (Life Technologies) and 500 mM DTT, separated in 4%–12% Bis-Tris SDS-polyacrylamide gels and transferred to a nitrocellulose membrane. The region corresponding to the protein size and up to the protein size plus 75 kDa was excised, and RNA was isolated following Proteinase K (NEB) treatment and column purification (Zymo). IP samples were subjected to dephosphorylation by alkaline phosphatase (FastAP, Thermo Fisher) and T4 PNK (NEB), followed by 3’ end adapter ligation (T4 RNA Ligase, NEB). After sequential low-salt and high-salt washes as in the standard eCLIP protocol, an additional wash with 2M urea buffer was performed to minimize nonspecific background. RNA was then isolated with direct Proteinase K (NEB) treatment and column purification (Zymo)^77^. Both IP and input RNA samples were then reverse transcribed and PCR-amplified according to the standard eCLIP workflow. Libraries were then sequenced on a NovaSeq X Plus (100 bp PE) platform.

#### eCLIP data analysis

eCLIP data was analyzed following the previously described pipeline^76^. Adapter trimmed (cutadapt v3.4) reads were mapped to the human genome using a custom STAR alignment pipeline as described above. PCR duplicates were identified and removed using UMI-tools^78^ Reads were also subjected to peak calling using Clipper^76^. Peaks were normalized to their paired input samples and subsequently annotated. High-confidence peaks were then identified based on filtering criteria of log_2_FC>=3 and -log10p>=3 and reproducibility across both biological replicates.

#### qPCR assays

AR17-primed cDNA from AR17 adapter-ligated RNA described above was amplified using Fast SYBR Green master mix (Thermo Fisher) with primers for RNAs of interest (Supplementary Table S1) on a QuantStudio Real-Time PCR system (Thermo Fisher). Relative levels were quantified using the ΔΔCt method^79^.

#### Western blots

Western blots were performed by separating proteins in SDS-polyacrylamide gels followed by transfer to nitrocellulose membranes using standard procedures. Membranes were incubated overnight at 4°C with mouse polyclonal anti-ZCCHC8 (Abcam, ab68739) at 1:500, rabbit polyclonal anti-MTR4 at 1:1,000 (Abcam, ab70551), rabbit anti-ZCCHC6 (TUT7) (Sigma, HPA020615) at 1:500, rabbit polyclonal anti-β-actin (Cell Signaling, #4967S) at 1:400 each in PBS with 0.1% Tween (PBST) and 5% nonfat milk. Secondary antibodies were HPR donkey anti-rabbit IgG (H+L) (Thermo Fisher, #31485), or HPR goat anti-mouse rabbit IgG (H+L) (Thermo Fisher, #31430) at 1:20,000 in PBST with 5% nonfat milk. Western blots were visualized using an Odyssey Fc imaging system (LI-COR).

#### Quantification and statistical analysis

Statistical tests were performed using python (v3.10.13) or R (v3.6.1). Data visualization was carried out using matplotlib (v3.8.3) or ggplot2 (v3.3.5) packages. For all experiments appropriate statistical tests were used as described in figure legends and the Method details section.

---

**Supplementary Figure S1 related to Figure 1.**
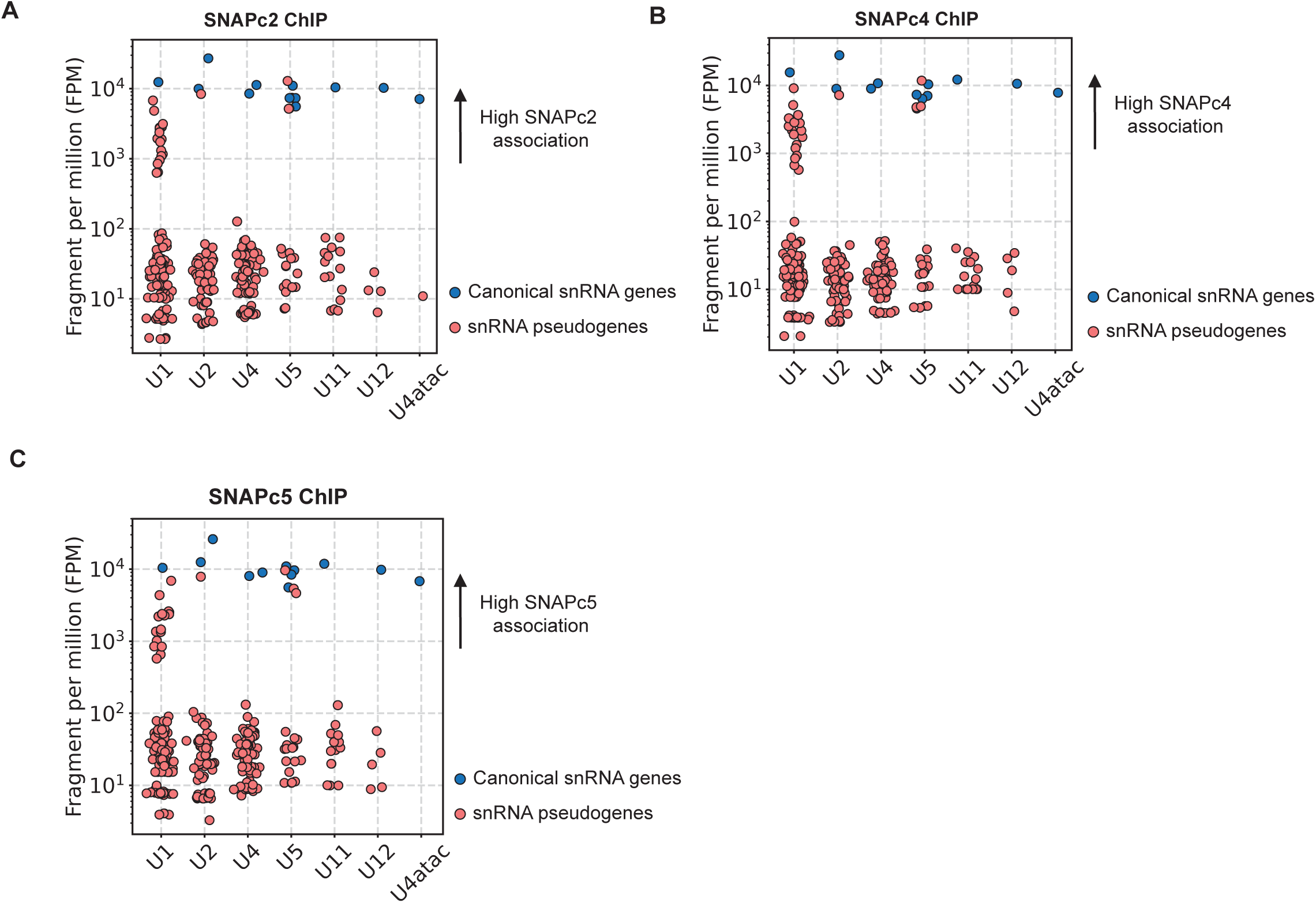
(A-C) Scatter plots of SNAPc2 (A), SNAPc4 (B), and SNAPc5 (C) ChIP seq reads, represented as fragments per million (FPM), for individual Pol II snRNA canonical genes (blue) and pseudogenes (red). Dataset from^38^.

**Supplementary Figure S2 related to Figure 3.**
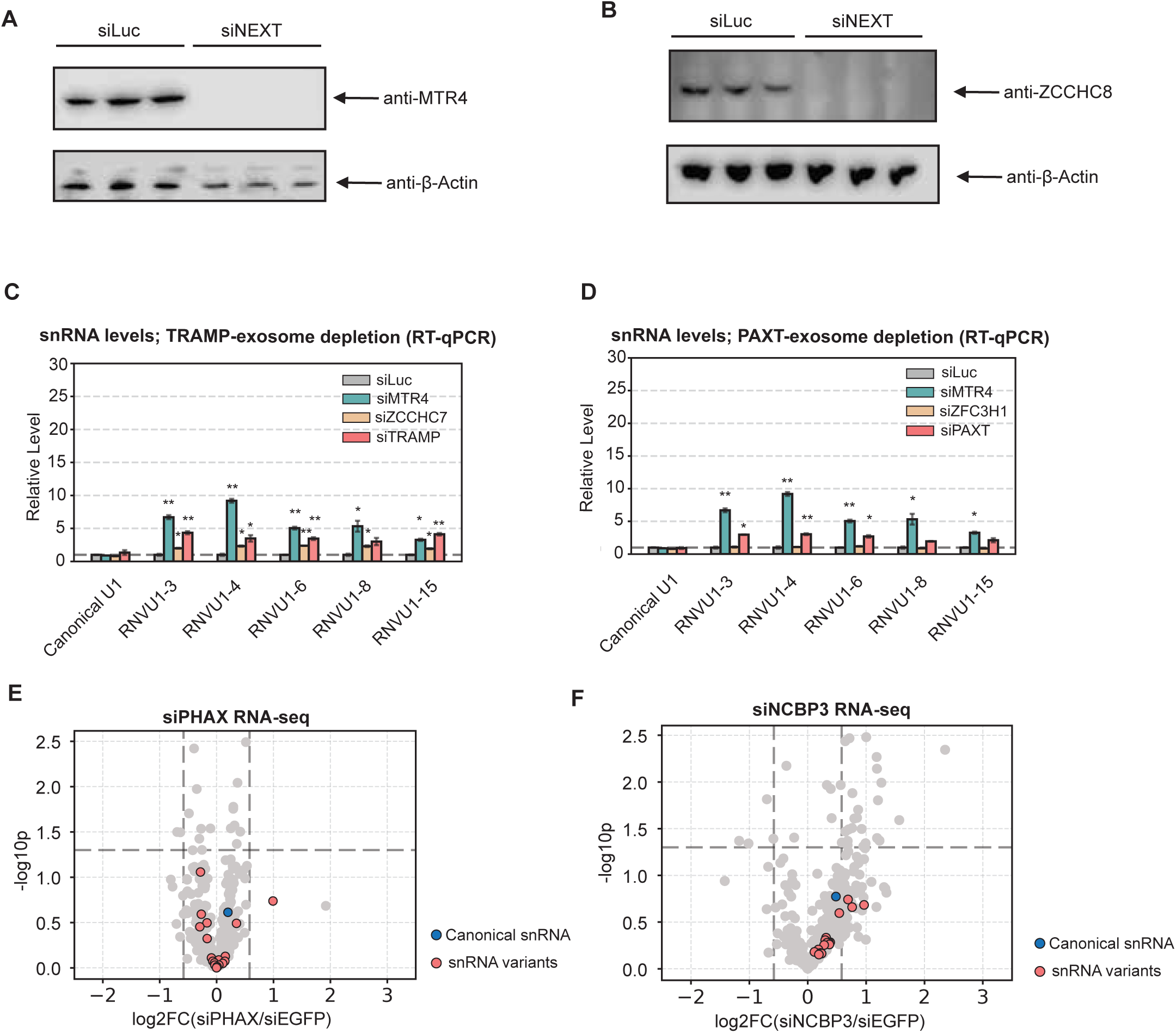
(A, B) Western blot showing depletion of MTR4 (A) and ZCCHC8 (B). β-tubulin serves as a loading control. (C) Relative levels of canonical U1 and U1 variants snRNAs upon depletion of TRAMP-exosome components (siTRAMP represents co-depletion of MTR4 and ZCCHC7) quantified by RT-qPCR and normalized to siLuc control conditions. P-values were calculated through two-tailed Student’s t-test (*: p<0.05, **: p<0.01). (D) Same as in (C) but under PAXT-exosome component depleted conditions (siPAXT represents co-depletion of MTR4 and ZFC3H1). (E, F) Volcano plots monitoring changes in sncRNA TPMs upon PHAX depletion (E) and NCBP3 depletion (F) relative to control depletion conditions monitored by RNA seq. P-values were calculated from three individual biological repeats by Student’s two-tailed t-test. Canonical snRNAs are labeled in blue, snRNA variants in red, and other sncRNAs in grey. Dataset is from^52^.

**Supplementary Figure S3 related to Figure 4.**
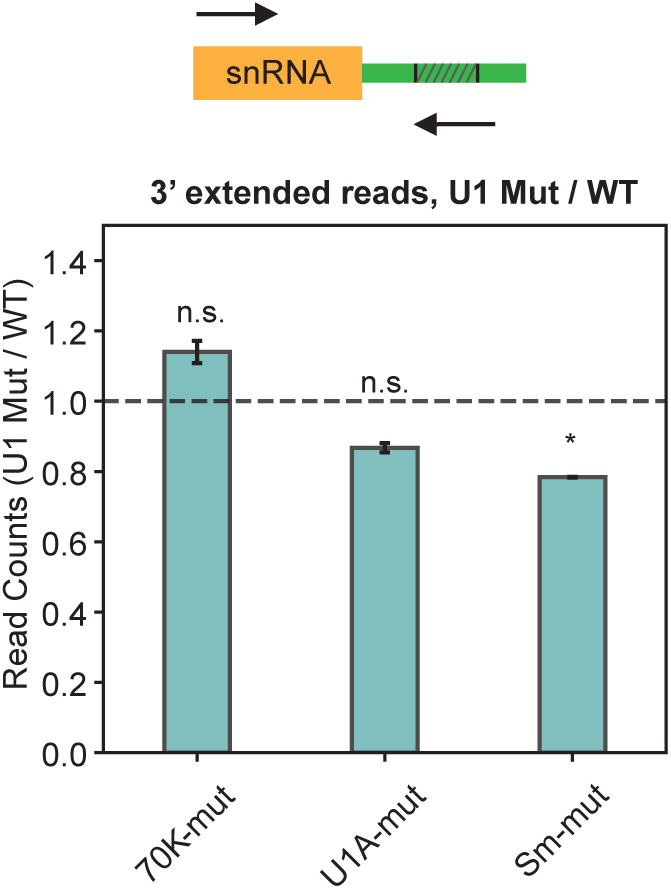
Relative levels of 3’ extended 70K-mut, U1A-mut, and Sm-mut U1 snRNAs, relative to co-expressed U1 WT, quantified by targeted sequencing. P-values are calculated by two-tailed Student’s t-test (n.s.: p>0.05, *: p<0.05).

**Supplementary Figure S4 related to Figure 5.**
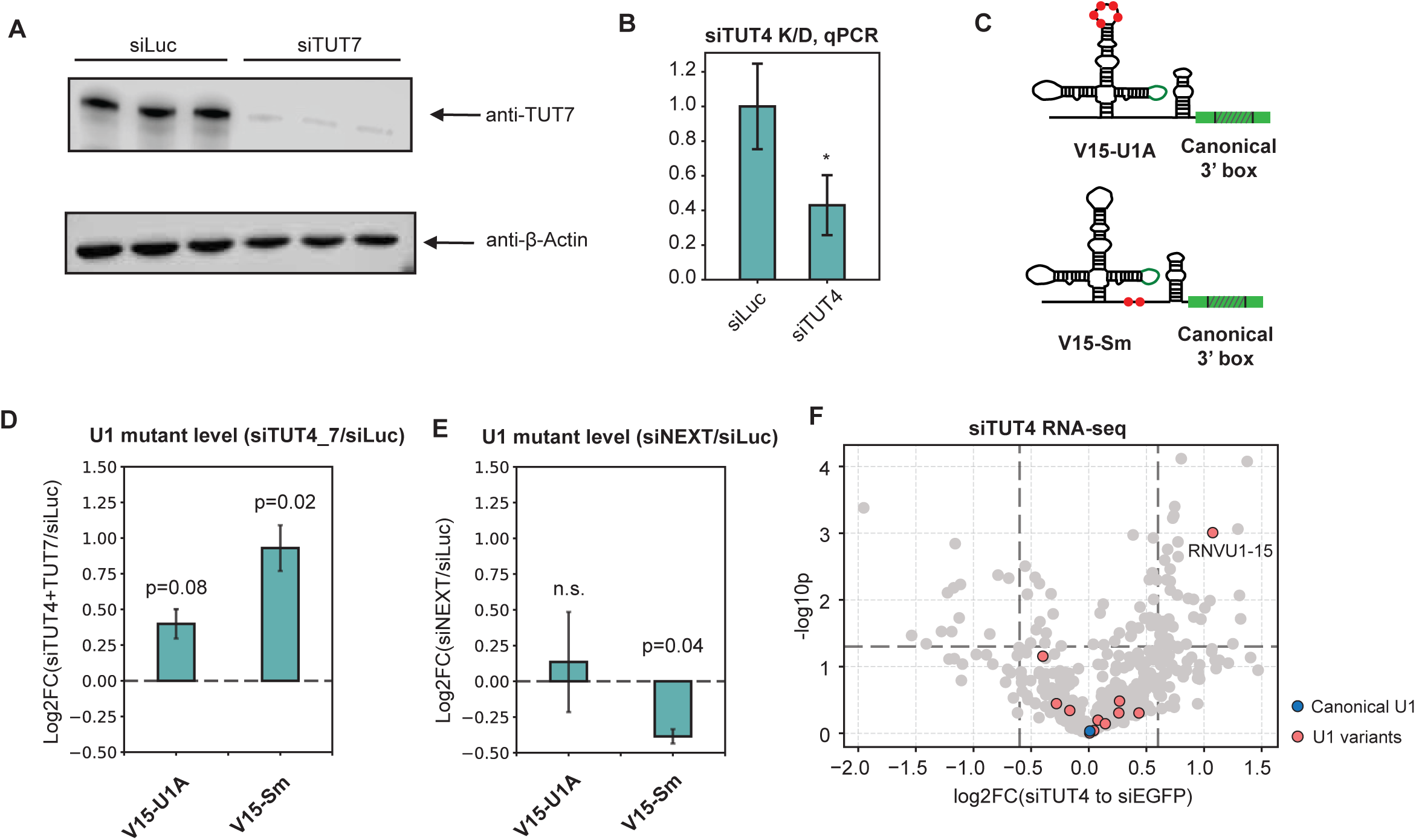
(A) Western blot showing depletion of TUT7. β-tubulin serves as a loading control. (B) Relative level of TUT4 mRNA in TUT4 depletion conditions compared to control (siLuc) conditions, measured by RT-qPCR. Samples were normalized to the level of Mitochondrial 12S rRNA. Error bars represent SEMs from three individual experiments; p-value was calculated by Student’s two-tailed t-test (*:p<0.1). (C) Schematics of U1 snRNA mutants with variant nucleotides from U1A (top) and Sm complex (bottom) binding sites of RNVU1-15. (D) Accumulation of V15 mutants in siTUT4/7 over siLuc control conditions, quantified by targeted RNA-seq. P-values are calculated by two-tailed Student’s t-test. (E) Same as (D) but comparing siNEXT to siLuc conditions. (F) Volcano plot showing changes in sncRNA accumulation in siTUT4 over siEGFP control conditions monitored by RNA-seq. P-values were calculated from three individual biological repeats by Student’s two-tailed t-test. Canonical snRNAs are labeled in blue, variant snRNAs in red, and other sncRNAs in grey. Dataset is from^58^.

**Supplementary Figure S5 related to Figure 6.**
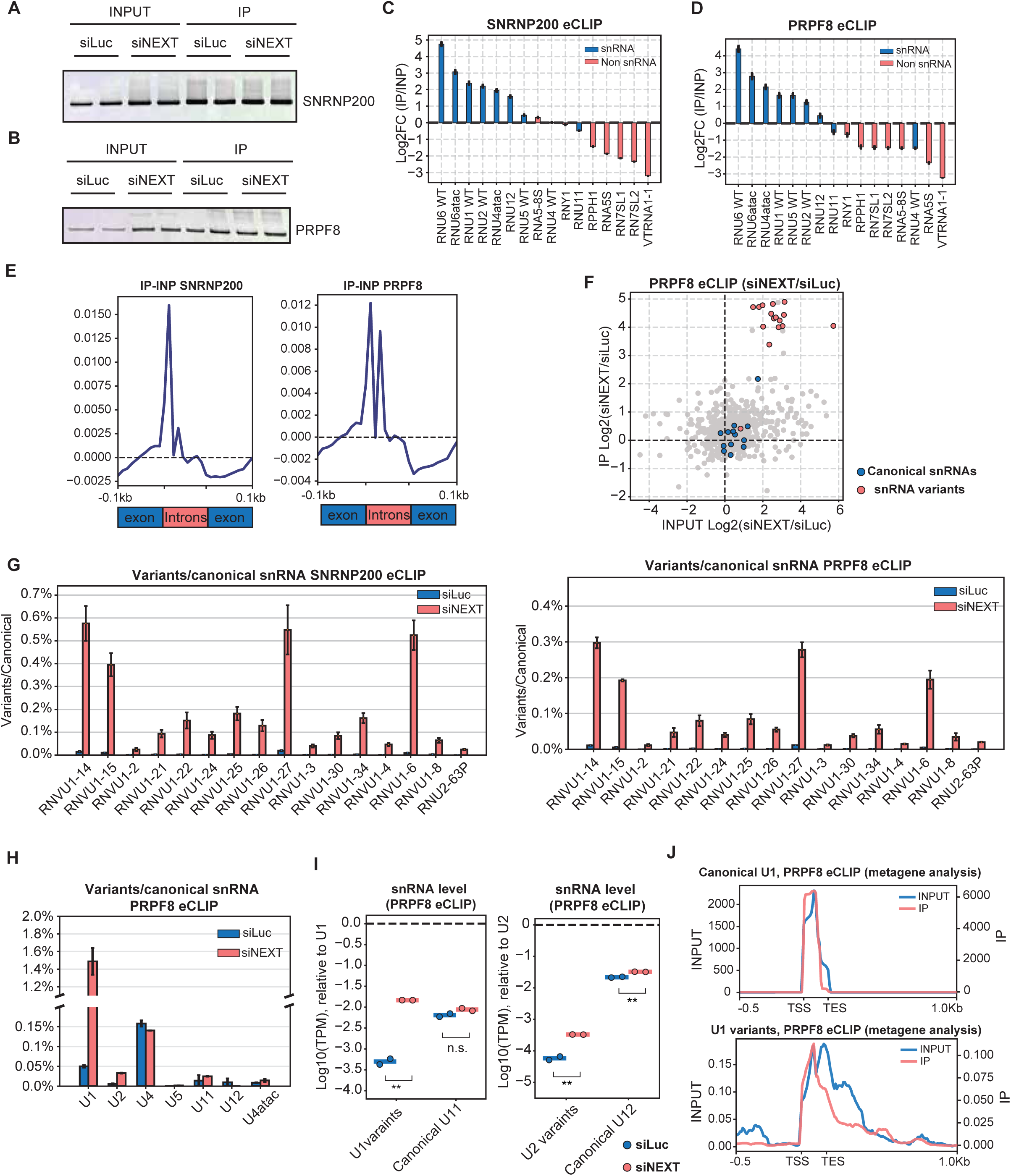
(A, B) Western blots for SNRNP200 (A) and PRPF8 (B) from eCLIP input and IP conditions. (C, D) Enrichment of snRNAs (blue) and representative other sncRNAs (red) in IP relative to input conditions in SNRNP200 (C) and PRPF8 eCLIP (D). (E) Metagene plot showing read distributions in eCLIP IP samples subtracted by read distributions in input samples over all annotated human introns and 0.1kb of flanking exons in aggregation. Introns are scaled to 0.1kb in length. (F) Scatter plot of small non-coding RNA enrichment in PRPF8 eCLIP, showing ratios of reads (measured as TPM) in siNEXT over siLuc conditions in Input versus IP fractions. Canonical snRNA are labeled in blue, U1 variants in red and other sncRNAs in grey. (G) Bar plot of ratios of snRNA variants relative to corresponding canonical snRNAs in SNRNP200 eCLIP (left) or PRPF8 (right) IP samples in siLuc (blue) or siNEXT (red) conditions. (H) Bar plot of the cumulative ratio of snRNA variants relative to corresponding canonical snRNAs in PRPF8 eCLIP IP samples in siLuc (blue) and siNEXT (red) conditions. (I) U1 variant (in aggregate) and canonical U11 snRNA levels relative to canonical U1 snRNA levels (left) (measured as TPM) or aggregate U2 variants and canonical U12 snRNA relative to canonical U2 snRNA (right) in PRPF8 eCLIP IP samples in siLuc (blue) and siNEXT (red) conditions. P-values are calculated through two-tailed Student’s t-test (n.s.: p>0.05, **: p<0.01). (J) Metagene plots showing read distributions for canonical U1 snRNA (upper panel) and U1 variants (lower panel) in PRPF8 eCLIP input (blue) and IP (red) conditions. Signals from 0.5Kb upstream of the transcription start site (TSS) to 1Kb downstream of the transcription end site (TES) are plotted.

**Supplementary Figure S6 related to Figure 6.**
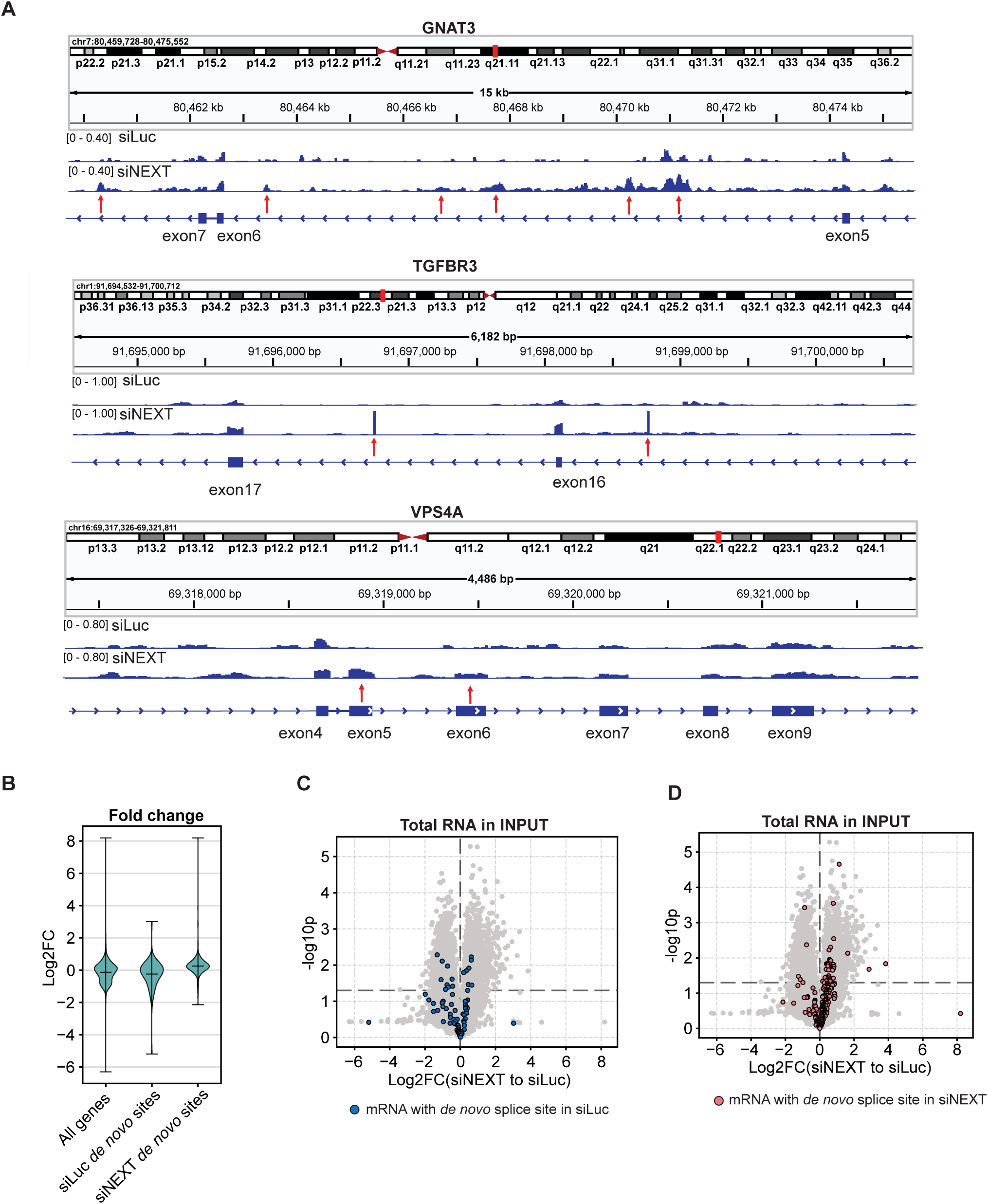
(A) Individual gene track plots representing altered splice site usage in siNEXT compared to siLuc conditions. Regions showing altered splicing are labelled by red arrows. (B) Violin plots showing relative level changes in siNEXT versus siLuc conditions of mRNA from all genes, genes with *de novo* splice sites in the siLuc condition, and genes with *de novo* splice sites in the siNEXT condition. (C,D) Differential expression in siNEXT over siLuc conditions of mRNAs from genes with *de novo* splice sites observed in siLuc conditions (C, blue) or in siNEXT conditions (D, red) (see Figure 6I), compared to all genes (grey).

**Supplementary Figure S7 related to Figure 7.**
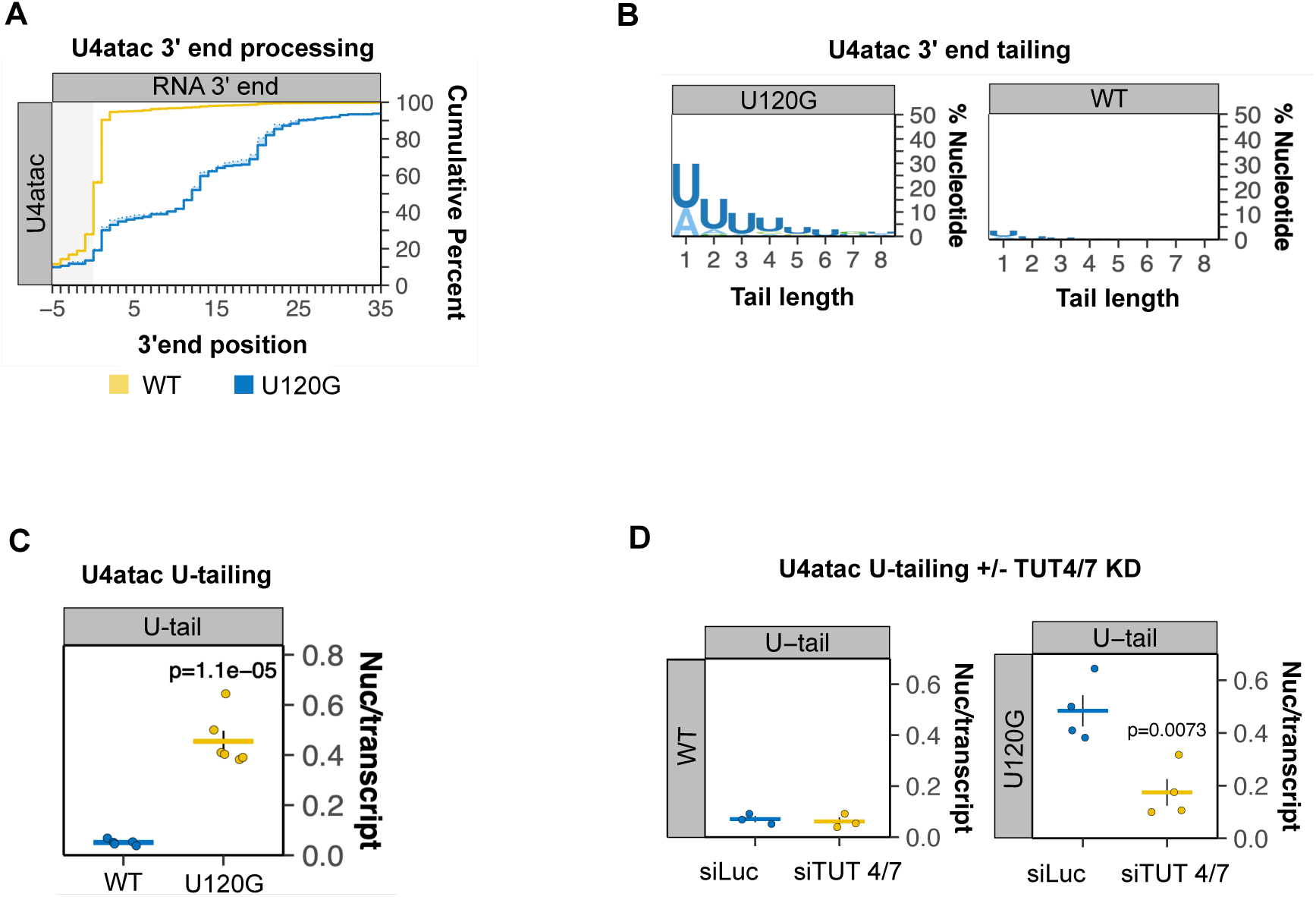
(A) Cumulative plot displaying cumulative fraction of 3’ end positions for U4atac WT (yellow) and U120G mutant (blue) snRNAs. (B) Logo plots showing post-transcriptional tail compositions of U4atac WT and U120G mutant snRNAs. (C) Average number of uridines observed in U4atac WT and U120G mutant snRNA post-transcriptional tails. (D) Average number of uridines observed in U4atac WT (left) and U120G mutant (right) snRNA post-transcriptional tails in TUT4/7 knockdown (siTUT4/7) and control knockdown (siLuc) conditions.

